# Presynaptic inhibition of cutaneous afferents prevents self-generated itch

**DOI:** 10.1101/806976

**Authors:** Augusto Escalante, Rüdiger Klein

## Abstract

Chronic itch represents an incapacitating burden on patients suffering a wide spectrum of diseases. Despite recent advances in our understanding of the cells and circuits implicated in the processing of itch information, chronic itch often presents itself without apparent cause. Here, we identify a spinal subpopulation of inhibitory neurons defined by the expression of Ptf1a involved in gating mechanosensory information self-generated during movement. These neurons receive tactile and motor input and establish presynaptic inhibitory contacts on mechanosensory afferents. Loss of Ptf1a neurons leads to increased hairy skin sensitivity and chronic itch, at least partially mediated through the classic itch pathway involving gastrin releasing peptide receptor (GRPR) spinal neurons. Conversely, chemogenetic activation of GRPR neurons elicits itch which is suppressed by concomitant activation of Ptf1a neurons. These findings shed new light on the circuit mechanisms implicated in chronic itch and open novel targets for therapy developments.

**Highlights:**

* Ptf1a specifies adult spinal presynaptic neurons contacting cutaneous afferents
* Loss of spinal Ptf1a+ neurons leads to self-generated itch and excessive grooming
* Absence of Ptf1a+ neurons increases hairy skin sensitivity which triggers scratching
* GRPR+ neurons act downstream of Ptf1a+ neurons in spontaneous itch

## Introduction

Itch is defined as an unpleasant sensation that elicits the desire or reflex to scratch (Hafenreffer, 1660). Despite its high prevalence, health and economic burden (Tripathi et al., 2019; Schut et al. 2019; Weisshaar et al., 2019) chronic itch remains a disease without cure and with very ineffective treatments to date (Oetjen et al., 2017). Under physiological conditions, chemical itch is initiated by receptors located in the skin, known as pruriceptors, which react to a plethora of chemical substances (Dong and Dong, 2018; Jakobsson et al., 2019). On the other hand, mechanical itch is triggered by light tactile stimuli, for example the sensation elicited by an insect crawling across the skin. It is also a widespread symptom associated with many skin and systemic diseases (Wahlgren et al., 1991; Yosipovitch and Bernhard, 2013; Mollanazar et al., 2016), but in healthy humans it has only been unequivocally induced under laboratory experimental conditions (Ikoma et al., 2005; Fukuoka et al., 2013) or under high intensity vibration (Mueller et al., 2018). Thus, identifying and understanding the neural circuits that mediate itch responses to mechanical stimuli remains a challenge.

The spinal cord dorsal horn is the first relay center for itch information. Excitatory spinal neurons expressing the gastrin releasing peptide receptor (GRPR) are required for itch sensation (Sun and Chen, 2007; Sun et al., 2009). Recently, evidence accumulated that only chemical but not mechanical itch signals are conveyed through the GRPR pathway. Excitatory spinal neurons expressing the neuropeptide Y1 receptor (Y1) specifically drive mechanical itch and are inhibited by neuropeptide Y (NPY)-expressing interneurons (Bourane 2015a; Acton et al., 2019). NPY neurons gate another population of excitatory spinal neurons marked by expression of Urocortin-3 (Ucn3). Reduced inhibition and increased excitation of Ucn3 neurons lead to chronic itch (Pan et al., 2019). Whether NPY neurons are the only inhibitory neurons that gate mechanical itch, is currently unclear. Moreover, the nature of the stimulus that induces spontaneous scratching and eventually results in self-inflicted skin wounds, has remained elusive.

The spinal dorsal horn is under strong inhibitory drive imposed by both local and supraspinal systems (Sorkin and Puig, 1996, 1998). The importance of maintaining a high level of inhibitory control over incoming somatosensory information in the spinal dorsal horn has long been recognized (Zeilhofer et al., 2012; Bardoni et al., 2013). Among inhibitory systems, spinal presynaptic inhibition plays a fundamental role in modulating various types of sensory afferents (Rudomin and Schmidt, 1999; Rudomin, 2009; Fink et al., 2014; Azim et al., 2014; Zimmerman et al., 2019). Inhibition at the terminals of sensory afferents allows for the reduction in excitability of downstream neurons without requiring postsynaptic inhibition of the target neurons themselves. Presynaptic inhibition is thought to reduce gain at sensory synapses and filter out irrelevant or self-generated sensory cues in order to preserve the ability to generate adequate responses to environmental stimuli in many animal species (Watson, 1992; Frost et al., 2003; Seki et al., 2003; Poulet and Hedwig, 2006). In the spinal dorsal horn, inhibitory interneurons named GABApre neurons, characterized by expression of Pancreas transcription factor 1a (Ptf1a), were identified as the source of presynaptic inhibitory synapses onto ventral proprioceptive and dorsal cutaneous afferents (Betley et al., 2009). GABApre neurons-mediated presynaptic inhibition was shown to control proprioceptive input to regulate fine motor tasks (Fink et al., 2014).

Dorsal horn inhibitory neurons share a common developmental origin. Spinal dorsal inhibitory neurons derive from the dI4/dILA class of interneurons defined during spinal cord neurogenesis through selective expression of defined transcription factor combinations (Gross et al., 2002; Müller et al., 2002; Helms and Johnson, 2003; Cheng et al., 2004; Cheng et al., 2005; Mizuguchi et al., 2006; Wildner et al., 2006). Ptf1a is transiently expressed in neural progenitors and acts as the main selector gene in establishing the dI4/dILA class (Glasgow et al., 2005). Specifically, Ptf1a is required for the specification of the subpopulations of inhibitory neurons implicated in itch processing: dynorphin (Huang et al., 2008; Wildner et al., 2013), glycinergic transporter 2 (Huang et al., 2008; Wildner et al., 2013; Borromeo et al., 2014) and neuropeptide Y (Huang et al., 2008; Bröhl et al., 2008; Wildner et al., 2013) positive neurons. According to these studies, there seems to be a common link between spinal inhibitory neurons responsible for presynaptic inhibition in the dorsal horn and subpopulations of inhibitory neurons demonstrated to play a role in the gating of somatosensory information whose loss leads to the development of chronic itch.

Here, we have used Ptf1a as a genetic entry point for manipulating these subpopulations and assessing their role in itch behavior. Our findings indicate that Ptf1a-derived inhibitory interneurons are essential components of a gating mechanism designed to block incoming self-generated mechanosensory information. Selective elimination of these neurons through intersectional genetics leads to the development of the highest frequency of spontaneous scratching behavior described to date. We find that other somatosensory modalities, including chemical itch, are generally not affected and gross motor behavior remained unchanged. These mice present increased hairy skin sensitivity and, following long-term video analysis in their own home cage, we show that the scratching phenotype is dependent on the level of activity of the animals. Finally, we show that GRPR neurons partially drive the scratching phenotype of Ptf1a neuron-ablated animals and that Ptf1a-derived interneurons control GRPR neuron-mediated itch. These results argue for Ptf1a as a common determinant of inhibitory neuron populations blocking innocuous mechanosensory input into the spinal dorsal horn. Moreover, they postulate chronic spontaneous scratching as a consequence of excessive entry of innocuous tactile information.

## Results

### Ptf1a-derived neurons comprise most of the inhibitory neurons in the dorsal horn

To begin testing the hypothesis that Ptf1a-derived neurons constitute a subpopulation of spinal inhibitory neurons gating mechanosensory information, we first analyzed the two existing Cre recombinase knock-in mouse lines available: Ptf1a-Cre^CVW^ (Kawaguchi et al., 2002) and Ptf1a-Cre^EX1^ (Nakhai et al., 2007). In the spinal dorsal horns of adult Ptf1a-Cre^CVW^ and Ptf1a-Cre^EX1^ mice carrying the Ai9^lsl-tdTomato^ reporter (Madisen et al., 2010), we found that Ptf1a-derived cells were 2-fold more abundant in the Ptf1a-Cre^EX1^ than in the Ptf1a-Cre^CVW^ mouse line (Figure S1A-E), possibly due to better recombination efficiency (Glasgow et al., 2005). Since Ptf1a-Cre^EX1^-derived cells recapitulated the same migration patterns previously described for Ptf1a-derived neurons in the Ptf1a-Cre^CVW^ mouse line (Glasgow et al., 2005; Meredith et al., 2009) (Figure S1F-J), we focused our research on the Ptf1a-Cre^EX1^ mouse line (referred to as Ptf1a-Cre from now on). To further characterize Ptf1a-derived neurons, we took advantage of the nuclear expression of β-galactosidase (β-gal) present in Tau^ds-DTR^ mice (Duan et al., 2014) (Figure 1A). β-gal positive, Ptf1a-derived neurons were found in the dorsal horn (Figure 1B-F), partially overlapping with Pax2 immunoreactive cells, a marker of inhibitory neurons in the spinal cord (Punnakkal et al., 2014) (Figure 1G). Partial overlap suggests the presence of Pax2 negative, Ptf1a-Cre positive neurons, or may result from incomplete recombination in the Ptf1a-Cre mice or incomplete immunolabeling with Pax2 antibodies. Further analysis during early postnatal stages revealed a similar overlap with Pax2, co-localization with Tfap2β, a known downstream target of Ptf1a (Wildner et al., 2013), and little co-expression with Zic2, a marker of excitatory dorsal horn neurons (Escalante et al., 2013; Paixão et al., 2019). Unexpectedly, we found co-localization with excitatory marker genes Lmx1b and Tlx3 (Figure 1H) (same results observed in Ptf1a-Cre^CVW^ mice, data not shown). These results suggest a change in transcriptional regulation of the Ptf1a locus in the Cre mouse lines.

**Figure 1.**
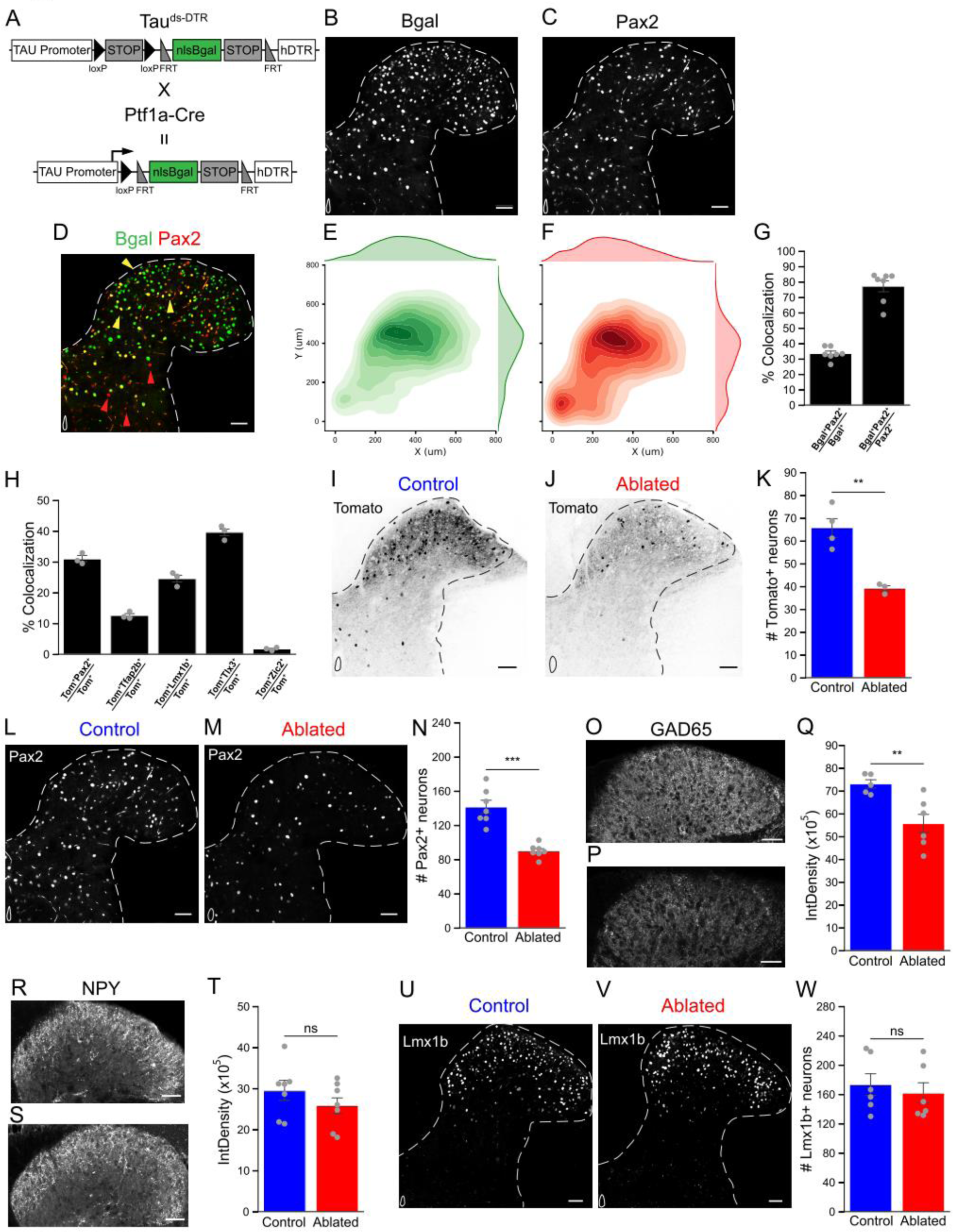
Ptf1a neurons are inhibitory and receive mechanosensory and motor input. **A.** Scheme representing the Cre/Flp intersectional Tau^ds-DTR^ mouse line used for diphtheria toxin (DT) mediated ablation. Nuclear β-galactosidase (βgal) expression tracks Cre^+^/Flp^−^ neurons. **B, C.** Representative images of βgal (**B**) and Pax2 (**C**) immunofluorescence of Ptf1a neurons in adult thoracic dorsal horn section. **D.** Merged image from panels B-C. Yellow arrowheads point to Pax2^+^βgal^+^ neurons and red arrowheads point to Pax2^+^βgal^−^. **E, F.** Kernel density estimate plots of the distribution of βgal^+^ (**E**) and Pax2^+^ (**F**) neurons. 88% of βgal^+^ neurons are located 200 µm dorsal to the central canal (8472 cells), similarly 75% of Pax2^+^ neurons are located dorsally (3680 cells), n=7 animals. **G.** Quantification of Pax2 and βgal co-localization in adult animals. Pax2^+^βgal^+^ double positive neurons represent 33.5±1.5% of βgal^+^ cells and 77.2±4% of Pax2^+^ cells, n=3-5 hemisections/animal, 7 animals. **H.** Characterization of Tomato^+^ neurons in postnatal day 0 spinal sections from Ptf1a-Cre; Ai9^lsl-tdTomato^ mice. Co-localization with Pax2 (30.9±1.03%), Tfap2β (12.6±0.67%), Lmx1b (24.55±1.33%), Tlx3 (39.6±1.35%) and Zic2 (1.72±0.3%); n=3 hemisections/animal, 3 animals. **I, J.** Representative images of adult thoracic spinal cord hemisection showing Ptf1a neurons expressing Tomato before (**I**) and one week after (**J**) DT administration. **K.** Quantification of the number of Tomato^+^ neurons in dorsal hemisections of control (65.8±4.5 cells) and Ptf1a neuron-ablated (39±1 cells) animals. n=6 hemisections/animal, 3-4 animals per group. Two-sided Student’s unpaired t test, (st=4.87, **p=0.0045). **L, M.** Representative images of Pax2 immunofluorescence in control (**L**) and Ptf1a neuron-ablated (**M**) dorsal hemisections. **N.** Quantification of the number of Pax2^+^ neurons in dorsal hemisections of control (141.36±7.75 cells) and Ptf1a neuron-ablated (89.96±2.92 cells) animals. n=3-4 hemisections/animal, 7 animals per group. Two-sided Student’s unpaired t test, (st=6.20, ***p=0.0003). **O, P.** Representative images of GAD65 immunofluorescence in control (**O**) and Ptf1a neuron-ablated (**P**) spinal cord dorsal horns. **Q.** Quantification of GAD65 immunofluorescence in the dorsal horns of control (7.3×10^6^±1.9×10^5^ AU) and Ptf1a neuron-ablated (5.5×10^6^±4.5×10^5^ AU) animals. n=10-12 hemisections/animal, 5-6 animals per group. Two-sided Student’s unpaired t test, (st=3.278, **p= 0.0095). **R, S.** Representative images of NPY immunofluorescence in control (**R**) and Ptf1a neuron-ablated (**S**) spinal dorsal horns. **T.** Quantification of Neuropeptide Y immunofluorescence in the dorsal horns of control (2.94×10^6^±2.44×10^5^ AU) and Ptf1a neuron-ablated (2.58×10^6^±3×10^5^ AU) animals. n=10-12 hemisections/animal, 7 animals per group. Two-sided Student’s unpaired t test, (st=1.10, ^ns^p=0.29). **U, V**. Representative images of Lmx1b immunofluorescence in control (**U**) and Ptf1a neuron-ablated (**V**) spinal dorsal horns. **W.** Quantification of the number of Lmx1b^+^ neurons in dorsal hemisections of the spinal cord in control (173.33±15.72 cells) and Ptf1a neuron-ablated (161.73±15.30 cells) animals. n=3-5 hemisections/animal, 6 animals per group. Two-sided Student’s unpaired t test, (st= 0.5287, ^ns^p= 0.6085). Scale bars: 50 µm. Data presented as mean ± SEM. See also Figure S1.

To further characterize Ptf1a neurons, we selectively ablated them in the adult spinal cord using an intersectional genetic approach. Combination of Ptf1a-Cre mice with a mouse line expressing Flp recombinase caudal to cervical region 5 (C5) during embryogenesis, Cdx2^FlpO^ (Britz et al., 2015), and the Tau^ds-DTR^ line allowed restricted expression of the human Diphtheria toxin receptor (DTR) in Ptf1a spinal neurons (Figure S1K-N). Neuronal ablation was achieved through the administration of diphtheria toxin (DT) at 7-8 weeks of age (Figure S1O). Based on co-expression of the intersectional Ai65D^ds-tdTomato^ reporter (Madisen et al., 2015), 40% of Ptf1a neurons were ablated in the thoracic spinal cord compared to controls treated with DT (Figure 1I-K). Numbers of Pax2^+^ dorsal spinal neurons were reduced by 36% (Figure 1L-N) in Ptf1a neuron-ablated mice, suggesting that approximately half of the 77% of Pax2 neurons derived from the Ptf1a lineage (see Figure 1G), were ablated. Consistently, GAD65, the marker of GABApre presynaptic terminals in the spinal dorsal horn (Hughes et al., 2005; Betley et al., 2009; Mende et al., 2016) decreased 24% (Figure 1O-Q). Markers of specific inhibitory neuron populations were unaffected in Ptf1a neuron-ablated mice, including Enkephalin, Galanin and nNOS (Figure S1P-R). Surprisingly, we did not detect a reduction in NPY (Figure 1R-T), suggesting that Ptf1a-Cre targeted a subset of inhibitory neurons, as suggested by Pax2 countings. The numbers of Lmx1b+ excitatory dorsal horn neurons were unchanged in Ptf1a neuron-ablated animals (Figure 1U-W). Such ablation bias has previously been reported using intersectional approaches (Duan et al., 2014). Sensory terminals of non-peptidergic IB4+ nociceptors, vGlut1+ Aβ and Aδ fibers, and vGlut2+ excitatory terminals were unaltered in Ptf1a neuron-ablated mice (Figure S1S-U).

### Ptf1a presynaptic inhibitory neurons receive mechanosensory and motor input

The Ptf1a lineage gives rise to presynaptic inhibitory neurons, named GABApre neurons (Betley et al., 2009; Fink et al., 2014; Mende et al., 2016). By expressing the Synaptophysin-GFP (SynGFP) fluorescent presynaptic marker in Ptf1a-Cre^+^ neurons (Tripodi et al., 2011) (Figure 2A), we found that 84% of presynaptic inhibitory contacts on adult vGlut1^+^ sensory terminals derive from Ptf1a neurons (Figure 2B-D), similar to early postnatal stages reported previously (Betley et al., 2009). After DT-mediated ablation of Ptf1a-derived neurons, we measured an 18% decrease in the numbers of presynaptic inhibitory vGAT^+^ contacts over vGlut1^+^ terminals compared to controls in the dorsal horns (Figure 2F).

**Figure 2.**
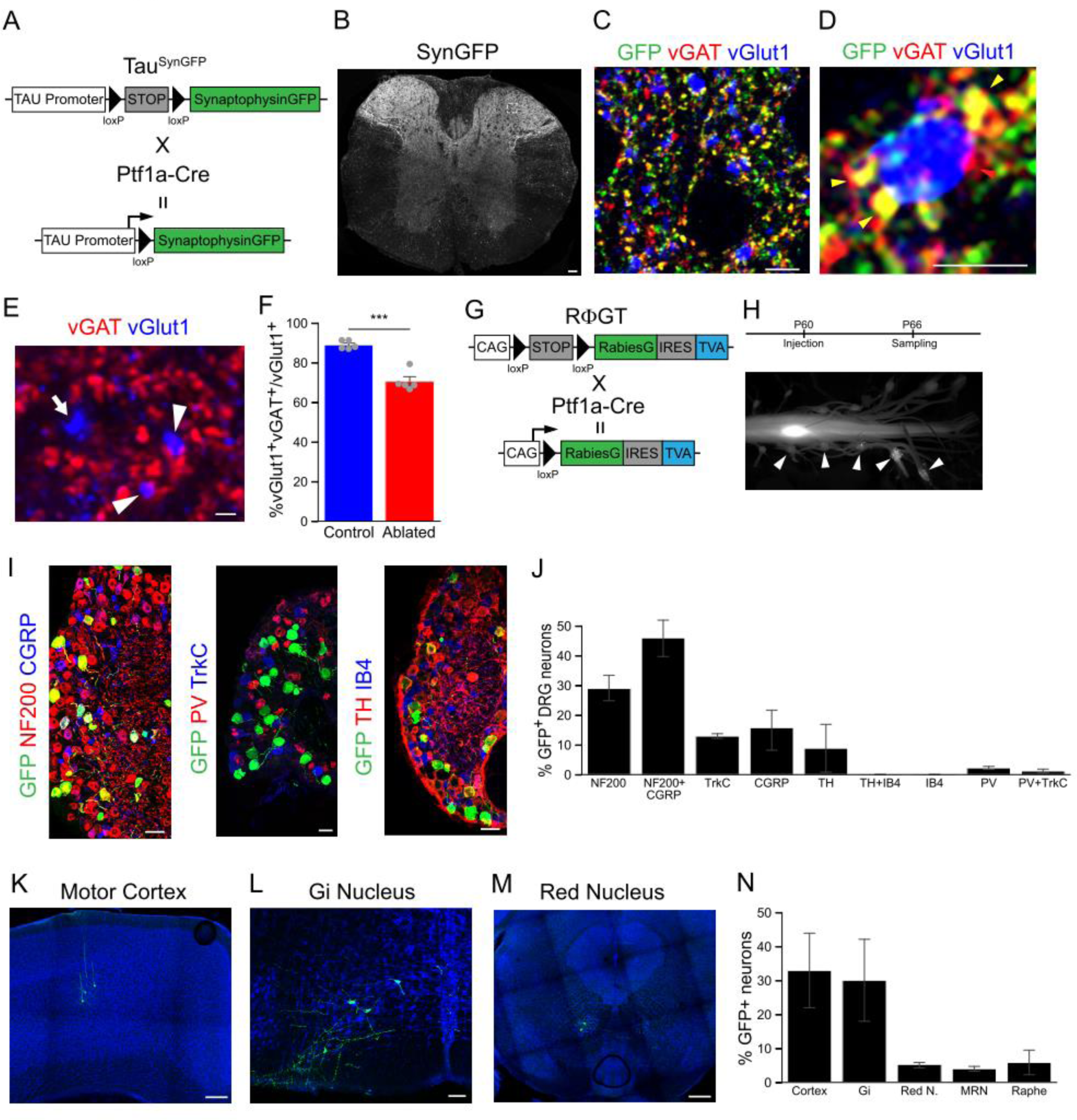
Loss of Ptf1a neurons leads to spontaneous scratching. **A.** Scheme representing the genetic strategy used for the labelling of Ptf1a presynaptic terminals. **B.** Overview of GFP immunofluorescence in thoracic spinal cord section from adult Ptf1a-Cre;Tau^lsl-SynGFP^ mouse. **C, D.** Representative image (**C**) of dorsal spinal cord showing sensory terminals (vGlut1^+^, blue) receiving presynaptic inhibitory (vGAT^+^, red) contacts. A high proportion are Ptf1a-derived GFP^+^ (green). High magnification image (**D**) showing a single sensory terminal expressing vGlut1^+^ (blue) contacted by several presynaptic inhibitory (vGAT^+^, red) terminals. Some derived from Ptf1a neurons (GFP^+^, yellow arrowheads) and one non-Ptf1a-derived vGAT^+^GFP^−^ terminal (red arrowhead). 84.36±2.66% of vGAT^+^ inhibitory terminals presynaptic to vGlut1^+^ sensory terminals are derived from Ptf1a neurons (n=3-4 hemisections/animal, 3 animals). **E.** Representative image of dorsal spinal cord showing sensory terminals (vGlut1^+^, blue) receiving presynaptic inhibitory (vGAT^+^, red) contacts (arrowheads). Arrow points to a single vGlut1^+^ sensory terminal devoid of presynaptic inhibitory contacts. **F.** Quantification of the percentage of vGlut1+ sensory terminals apposed by vGAT+ inhibitory synapses in the dorsal horn of control (blue, 88.99±0.96%) and Ptf1a neuron-ablated (red, 70.53±2.44%) animals. n=3 sections/animal, 5 animals/genotype. Two-sided Student’s unpaired t test, (st=7.5, ***p=0.00044). **G.** Scheme representing the RΦGT mouse line used for transsynaptic tracing with EnvA-pseudotyped Rabies virus. Cre recombination leads to the expression of Rabies G protein and TVA. **H.** Spinal cords, DRGs and brains were processed for immunofluorescence 6 days after injection. Arrowheads indicate DRGs with GFP^+^ sensory neurons. **I.** Representative images of DRGs showing GFP^+^ neurons labeled by transsynaptic tracing from Ptf1a neurons. Neurofilament (NF200), Calcitonin gene-related peptide (CGRP), Parvalbumin (PV), Tropomyosin receptor kinase C (TrkC), Tyrosine hydroxylase (TH), and Isolectin GS-IB4 (IB4) immunofluorescence. **J.** Quantification of GFP co-localization with sensory neuron markers expressed as percentage of the total of GFP^+^ neurons. n=3-5 sections/animal, 3 animals per marker combination. **K-M.** Representative images of GFP+ cortical (**K**), Gi nucleus (**L**) and red nucleus (**M**) neurons labelled by transsynaptic tracing from Ptf1a spinal neurons. **N.** Quantification of supraspinal input regions to Ptf1a spinal neurons expressed as percentage of the total of GFP^+^ neurons found in the brain. n=3-5 sections/animal, 5 animals. Scale bars: B, I: 50 µm; K, M: 200 µm; L: 100 µm; C: 5 µm; D: 2.5 µm; E: 1 µm. Data presented as mean ± SEM.

To gain insights into the monosynaptic inputs received by Ptf1a neurons, we performed retrograde tracing with EnvA-pseudotyped rabies virus (SADΔG-GFP) (Wickersham et al., 2006). Ptf1a-Cre mice were crossed to RΦGT (Takatoh et al, 2013) mice conditionally expressing the auxiliary TVA and G proteins (Figure 2G) directing specific infection of SADΔG-GFP to Ptf1a neurons. Injections were performed in adult animals and analyzed 6 days later. We found numerous GFP+ input neurons in 5-6 adjacent DRGs ipsilateral to the injection site (Figure 2H). Immunohistochemical characterization of GFP^+^ DRG neurons (Figure 2I-J) revealed low threshold mechanoreceptors (LTMRs) involved in transmission of touch information, marked by NF200^+^ (32.04±4.26%), NF200^+^CGRP^+^ (43.73±6.85%), TrkC^+^ (12.76±0.81%), and TH^+^ (8.86±8.04%). A small percentage of them were associated with unmyelinated peptidergic nociceptive neurons: CGRP^+^NF200^−^ (12.6±7.07), very few with proprioceptive inputs: PV^+^ (2.12±0.5%) and PV^+^TrkC^+^ (1.06±0.63) and none with non-peptidergic nociceptors (IB4^+^) (Lallemend and Ernfors, 2012; Bourane et al., 2015b; Usoskin et al., 2015). Ptf1a neurons also received monosynaptic inputs from supraspinal regions, most of which were associated with descending motor pathways (Figure 2K-M). Most prominent regions were layer V motor cortex (32.95±12%), corresponding to corticospinal projection neurons involved in skilled movements (Moreno-Lopez et al, 2016; Ueno et al., 2018), and the gigantocellular nucleus (Gi) (30.1±12.33%) of the reticulospinal tract important for motor control (Bouvier et al., 2015; Liang et al., 2016; Capelli et al., 2017; Brownstone and Chopek, 2018). Minor contributions came from the red nucleus, the medullary reticular nucleus, and the raphe nucleus (Figure 2N). In summary, these results revealed that Ptf1a neurons established presynaptic inhibitory contacts with myelinated cutaneous afferents and received direct mechanosensory and higher motor information.

### Ablation of Ptf1a spinal neurons leads to spontaneous scratching

To detect early, spontaneous behavioral alterations in Ptf1a neuron-ablated mice, we video recorded the mice daily in their home cages before and after DT treatment. Ptf1a neuron-ablated animals developed an intense itch sensation and started to scratch only 2 days after the end of DT administration (Day 5 since the beginning of the experiment) and reached peak scratching frequency just one day later (Figure 3A). At day 7, animals started to develop fur loss and skin injuries, so the experiment was terminated. Scratching was mostly directed to the nape of the neck, the forearm and flanks. Very rarely we observed scratching towards the cheeks, snout or scalp, in accordance with the limitation in spinal rostral ablation imposed by the use of Cdx2^FlpO^ which spares segments receiving somatosensory information from those dermatomes (Takahashi et al., 1992). In summary, Ptf1a neuron-ablated animals scratched more than 60-fold compared to control littermates after DT treatment (Figure 3B). This behavior developed very quickly, reaching peak values 6.8 days after first DT administration (Figure 3C). We also quantified grooming time and found that Ptf1a neuron-ablated animals spent 5-fold more time grooming than their control littermates after DT treatment (Figure 3D). To further analyze the behavioral consequences of Ptf1a neurons-ablation, we performed a battery of somatosensory behavioral assays. The spontaneous scratching phenotype was not related to chemical itch, as injections of chemical histaminergic and non-histaminergic pruritogens in the back region of Ptf1a neuron-ablated animals before the onset of spontaneous scratching, but after loss of Ptf1a neurons (Figure S2A), showed no differences compared to control littermates (Figure 3E-F). Additional behavioral tests revealed no changes in the processing of other somatosensory modalities (Figure S2B-F), acute mechanical itch (Figure S2G-H), locomotor coordination (Figure S2I-J) and anxiety (Figure S2J). Taken together, these findings suggest that the subset of Ptf1a^+^ inhibitory neurons that are targeted by Ptf1a-Cre, specifically mediate mechanical itch control in the spinal dorsal horn.

**Figure 3.**
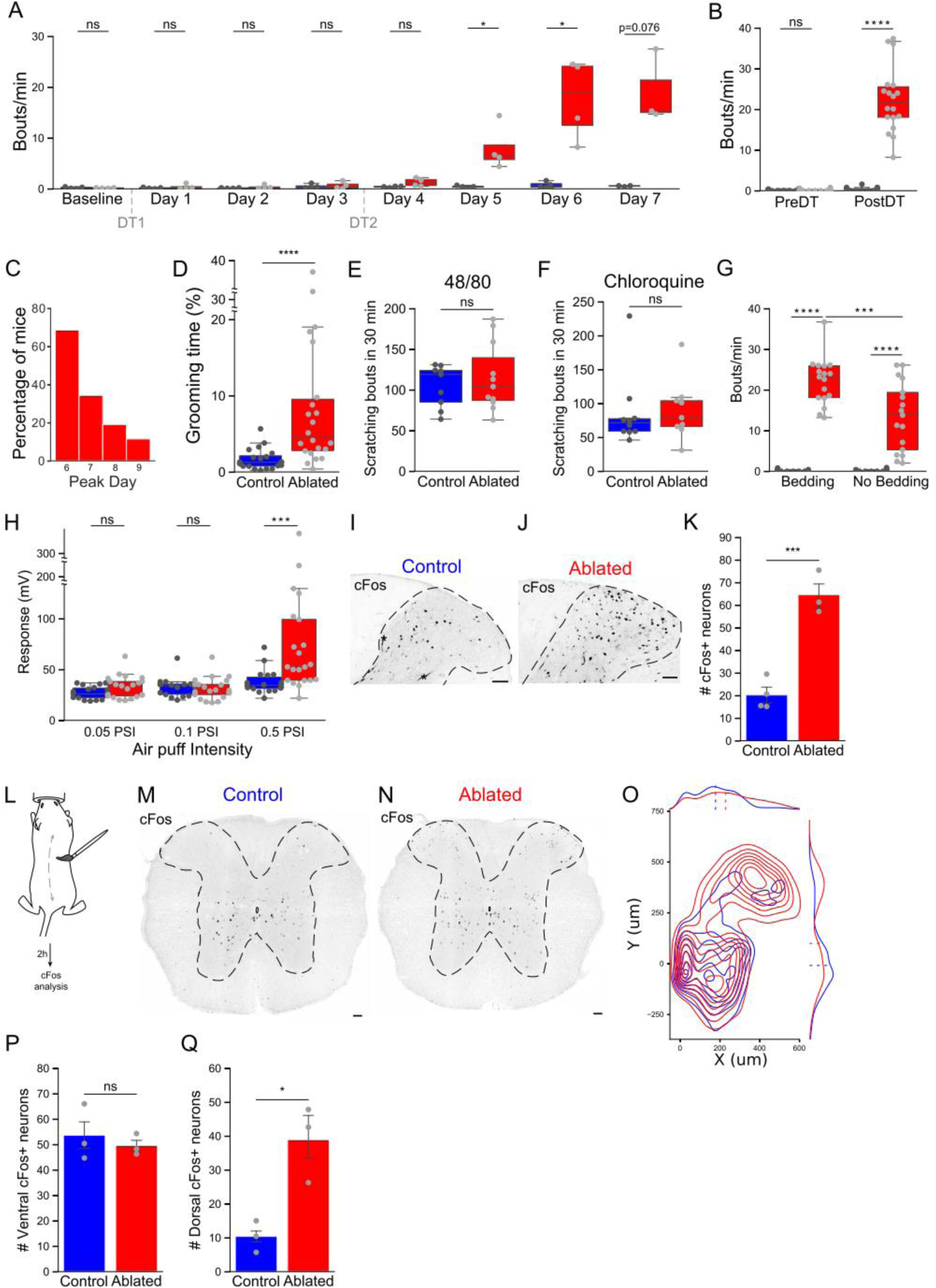
Ptf1a neurons control hairy skin sensitivity. **A.** Time course of the spontaneous scratching phenotype in control (blue) and Ptf1a neuron-ablated (red) animals. DT1 and DT2 depicts the times of DT administration. n=4 animals per group and time point. Day 7 only includes 3 animals due to the sacrifice of one animal according to ethical considerations. Two-sided Mann Whitney’s unpaired u test, ^ns^p>0.05, *p<0.05. **B.** Quantification of scratching frequency in control (blue) and Ptf1a neuron-ablated (red) animals in home cages, before and after DT treatment. n =16/18 animals per group. Control: PreDT: 0.187±0.044 vs PostDT: 0.337±0.111 bouts/minute; Ablated: PreDT: 0.166±0.050 vs PostDT: 22.40±1.918 bouts/minute. Two-sided Mann Whitney’s unpaired u test: PreDT (st= 159.5, ^ns^p= 0.5944), PostDT (st=0, ****p= 6.79×10^−7^). **C.** Histogram representing percentage of Ptf1a neuron-ablated animals at maximum scratching frequency in relation to time after DT1: Day 6 68.57%, Day 7 34.28%, Day 8 19.04%, Day 9 11.4%. n = 35 animals. **D.** Quantification of time spent grooming for control (blue) and Ptf1a neuron-ablated (red) animals in home cages after DT treatment. n =20-22 animals per group. Control: 1.7±0.31%, Ablated: 8.97±2.13%. Two-sided Mann Whitney’s unpaired u test, (st=65.0, ****p=9.98 x10^−5^). **E, F.** Quantification of scratching frequency in control (blue) and Ptf1a neuron-ablated (red) animals after injection of compound 48/80 (**E**) or chloroquine (**F**) in the back 5 days after DT1. n=9-11 animals per group. **E.** Control: 105.11±8.62 bouts/minute, Ablated: 115.27±12.64 bouts/minute. Two-sided Mann Whitney’s unpaired u test, (st=48.5, ^ns^p=0.97). **F.** Control: 85.5±16.78 bouts/minute, n=10 mice, Ablated: 89.77±14.66 bouts/minute, n=9 mice. Two-sided Mann Whitney’s unpaired u test, (st=35.0, ^ns^p=0.4377). **G.** Quantification of scratching frequency in control (blue) and Ptf1a neuron-ablated (red) animals in home cages 30 minutes after removal of bedding material. n = 14-17 animals per group. Control: Bedding: 0.17±0.06 vs No Bedding: 0.17± 0.07 bouts/minute; Ablated: Bedding: 22.16±1.4 vs No Bedding: 13.43±2.04 bouts/minute. Two-sided Mann Whitney unpaired u test, Bedding (st= 0, ****p= 2.31×10^−6^), No Bedding (st=0, ****p= 2.207×10^−6^). Two-sided Wilcoxon’s signed-rank paired test, Ablated (st=4.0, ***p=0.0006). **H.** Quantification of the activity response elicited by air puffs of increasing intensity directed towards the back of control (blue) and Ptf1a neuron-ablated (red) animals 5 days after DT1. n = 12-21 animals per group. 0.05PSI: Control 27.25±1.46 vs Ablated 32.8±2.4 mV, 0.1PSI: Control 32.41±2.2 vs Ablated 31.4±2.3 mV, 0.5PSI: Control 37.7±2.8 vs Ablated 81.4±15.3 mV. Two-sided Mann Whitney’s unpaired u test, ^ns^p>0.05, ***p < 0.001. **I, J.** Representative images of cFos immunofluorescence in control (**I**) and Ptf1a neuron-ablated (**J**) spinal cord dorsal hemisections after DT treatment. Asterisks: example of non-specifically labelled blood vessels. **K.** Quantification of the number of cFos^+^ neurons in dorsal hemisections in control (20.2±3.34 cells) and Ptf1a neuron-ablated (64.66±5.55 cells) animals. n=6-8 hemisections/animal, 3-4 animals per group. Two-sided Student’s unpaired t test, (st=−7.289, ***p=0.00076). **L.** Schematic representation of cFos induction through fur stimulation. **M, N.** Representative images of cFos immunofluorescence in control (**M**) and Ptf1a neuron-ablated (**N**) spinal cord sections after DT treatment and fur stimulation protocol. Notice how the number of cFos^+^ neurons changes in dorsal, but not ventral horns. **O.** Kernel density estimate plot of the distribution of cFos^+^ neurons in control (blue) and Ptf1a neuron-ablated (red) spinal cords after fur stimulation. Ptf1a neuron-ablated animals show increased number of cFos^+^ dorsal neurons compared to control animals. Control: n=3 animals, 3075 neurons; Ablated: n= 3 animals, 3957 neurons. **P, Q.** Quantification of the number of cFos^+^ neurons in ventral (**P**) and dorsal (**Q**) spinal cord from control (blue) and Ptf1a neuron-ablated (red) animals after fur stimulation. n=6-8 hemisections/animal, 3 animals per group. **P.** Ventral: Control 53.7±6.4 vs Ablated 49.6±2.43 cells, two-sided Student’s unpaired t test, (st=0.60, ^ns^p=0.58). **Q.** Dorsal: Control 10.4±2.7 vs Ablated 38.9±6.5 cells, two-sided Student’s unpaired t test, (st=−4.056, *p=0.015). Scale bars: 50 µm. Data presented as mean ± SEM. See also Figure S2.

### Ptf1a neurons control hairy skin sensitivity

To identify the stimuli that induce itch in Ptf1a neuron-ablated mice, we studied the scratch response of mice at peak scratching phase before and after removal of bedding material from the home cage. Ptf1a neuron-ablated animals decreased 40% in scratching frequency (Figure 3G) suggesting a role for environmental stimuli in the induction of itch. We reasoned that mechanosensory stimuli causing the scratch phenotype might be detected by hairs, the most common mechanoreceptor in hairy skin, and performed an experiment to test hairy skin sensitivity to air puffs of different intensities directed towards the back of the animal (Orefice et al., 2016). First, we evaluated the response curve to air puffs of increasing intensities for control animals treated with DT (Figure S2K). The minimum intensity that elicited a significant response was 0.5 PSI. Ptf1a neuron-ablated animals showed a 2-fold increased response to this threshold intensity compared to controls (Control 37.71±2.81 vs Ablated 81.4±15.27 mV) (Figure 3H). Ablation of Ptf1a neurons did not alter startle responses or pre-pulse inhibition characteristics in both the acoustic and tactile versions of the assay (Figure S2L-Q).

According to these findings, ablation of Ptf1a neurons leads to heightened hairy skin sensitivity and intense spontaneous scratching that depends on the abundance of mechanosensory cues in the environment. Loss of Ptf1a neurons leads to increased activity in the dorsal horns as measured by the number of cFos^+^ neurons (Figure 3I-K). In an attempt to establish a causal relationship between hairy skin sensitivity and dorsal horn neuronal activity and at the same time avoiding the potential effect of scratching in dorsal horn activity, we designed a back hair stimulation protocol under anesthesia (Figure 3L). Spinal cords of Ptf1a neuron-ablated animals whose back hair had been stimulated, showed a localized increase of cFos^+^ neurons in the dorsal horns almost 4-fold higher than in control animals (Figure 3M-O). As an internal control, the number of cFos^+^ neurons in the ventral spinal cord, a region not innervated by cutaneous afferents, was not altered (Figure 3P-Q). These results support the idea that the spontaneous scratching phenotype observed in Ptf1a neuron-ablated animals is a response to hair self-stimulation that is inhibited by Ptf1a neurons under physiological conditions.

### Mouse activity correlates with scratching in Ptf1a neuron-ablated animals

In the pursuit of understanding the scratching behavior of Ptf1a-ablated animals in as naturalistic as possible environment for laboratory mice (Babayan and Konen, 2019), we designed and built a system to video record individual mice in their own home cages continuously for several days without human interaction (Figure 4A). A combination of machine learning methods was used to extract the frames in which the animal performed scratching behavior, and to analyze the resulting data (Figure S3A-D). Our aim was to understand if there were differences in the intensity of the phenotype depending on the circadian phase and how we could explain the association between hairy skin and scratching.

**Figure 4.**
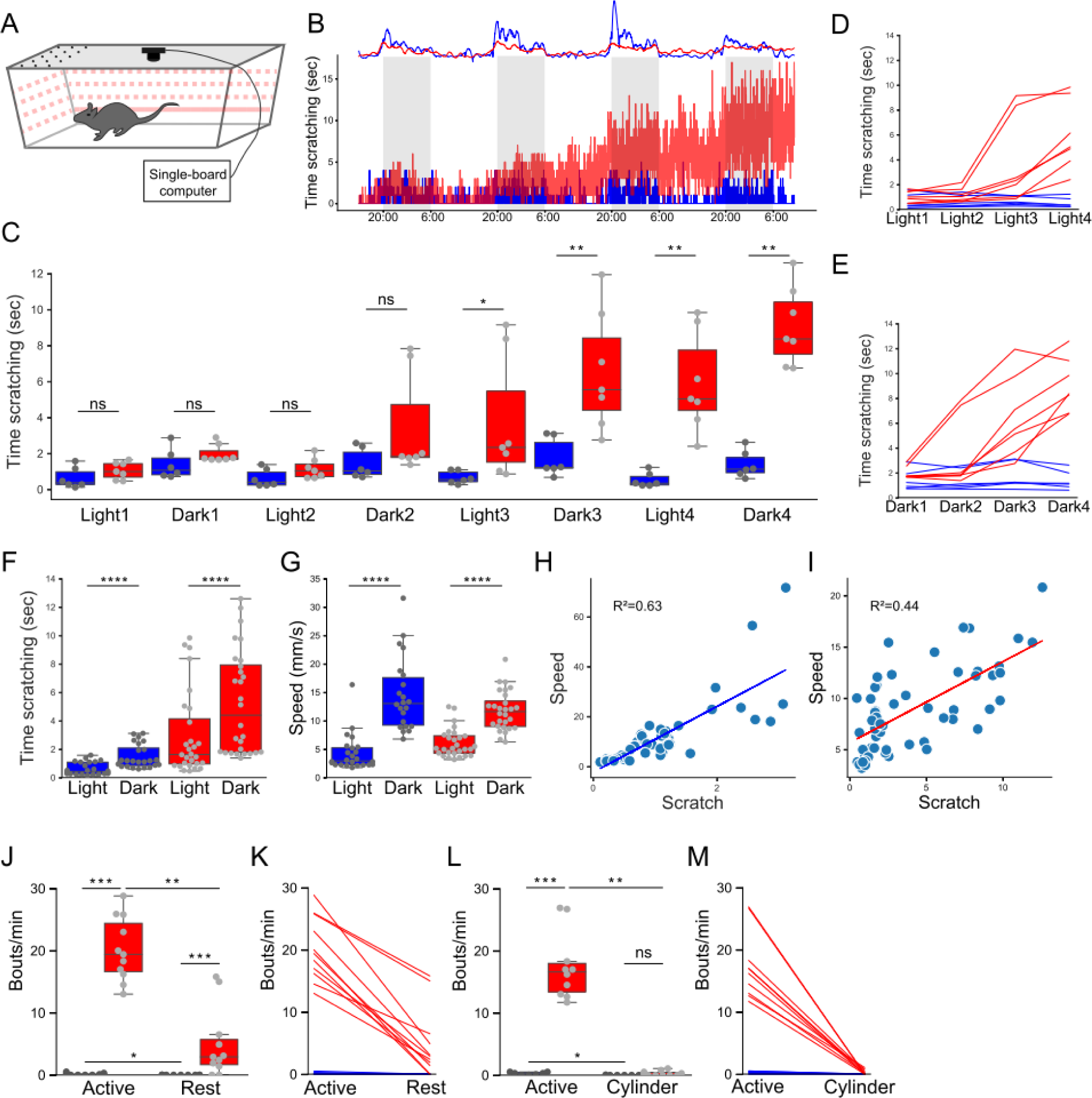
Scratching in Ptf1a neuron-ablated animals depends on animal activity. **A.** Schematic representation of the setup used to record the activity of single housed mice continuously for several days (see Materials and Methods for details). **B.** Graphical representation of the time spent scratching as a function of time. Top panel represents average speed for control (blue, total average speed: 10.6±3.2 mm/s, n=6 mice) and Ptf1a neuron-ablated (red, total average speed: 8.74±0.97 mm/s, n=7 mice) animals, two-sided Mann Whitney’s unpaired u test: (st=19.0, p=0.83). Bottom panel represents the average scratching time per minute for control (blue, total average time: 0.98±0.25 seconds, n=6 mice) and for Ptf1a neuron-ablated (red, total average time: 4.48±0.78 seconds, n=7 mice), two-sided Mann Whitney’s unpaired u test: (st=0.0, **p=0.0034). Gray shadows represent the dark phase of the light cycle. **C.** Scratching time per minute in each light/dark phases for control (blue, n=6 mice) and Ptf1a neuron-ablated (red, n=7mice) animals during the entire length of the video recordings. Two-sided Mann Whitney’s unpaired u test: ^ns^p>0.05, *p<0.05, **p<0.01. **D, E.** Time course change of mean scratching time per minute in each light (**D**) and dark (**E**) phase for each control (blue, n=6 mice) and Ptf1a neuron-ablated (red, n=7 mice) animals. Each Ptf1a neuron-ablated animal shows higher scratching during the dark than during the corresponding light phase. **F.** Mean scratching time per minute in control (blue, Light: 0.61±0.09 seconds, Dark: 1.48±0.17 seconds, n=6 mice) and Ptf1a neuron-ablated (red, Light: 2.97± 0.56 seconds, Dark: 5.26± 0.68 seconds, n=7 mice) animals during light and dark phases. Two-sided Wilcoxon’s signed-rank paired test, Control Light vs Dark (st=0.0, ****p=1.94×10^−5^), Ptf1a neuron-ablated Light vs Dark (st=0.0, ****p=4 x10^−6^). **G.** Mean speed in control (blue, Light: 4.3±0.65 mm/s, Dark: 14.55±1.37 mm/s, n=6 mice) and Ptf1a neuron-ablated (red, Light: 6.1±0.47 mm/s, Dark: 11.73±0.65 mm/s, n=7 mice) animals during light and dark phases. Two-sided Wilcoxon’s signed-rank paired test, Control Light vs Dark (st=−4.1, ****p=4.01 x10^−5^), Ptf1a neuron-ablated Light vs Dark (st= −4.6, ****p=4.23 x10^−6^). **H, I.** Ordinary linear regression analysis model fitting for scratching and speed data of control (**H**) and Ptf1a neuron-ablated (**I**) animals during each cycle phase for the whole study. Control [F(1,46)=79.54, p=1.36×10^−10^)], n=6 animals; Ablated [F(1,54)=43.12, p=2.08×10^−8^)], n=7 animals. Note: Y axis scale in H different from I. **J.** Scratching frequency in control (blue) and Ptf1a neuron-ablated (red) animals at peak day in home cages (Active) and 30 minutes after returning them to their respective racks (Rest). Control: active (0.12±0.05 bouts/min), rest (0±0 bouts/min), n=10 mice; Ablated: active (20.17±1.54 bouts/min), rest (4.9±1.67 bouts/min), n=11 mice. Two-sided Mann Whitney’s unpaired u test, Active: (st=0.0, ***p=0.0001), Rest: (st=10.0, ***p=0.0005). Two-sided Wilcoxon’s signed-rank paired test, Control: (st=0.0, *p=0.031), Ablated: (st=0.0, **p=0.003). **K.** Individual mice representation of data in G for Control (blue lines) and Ptf1a neuron-ablated (red lines) animals. **L.** Scratching frequency in control (blue) and Ptf1a neuron-ablated (red) animals at peak day in their home cages (Active) and 1 hour later inside the startle response cylinder (Cylinder). Control: active (0.243±0.08 bouts/min), cylinder (0.013±0.01 bouts/min), n=7 mice; Ablated: active (17.39±1.71 bouts/min), cylinder (0.281±0.12 bouts/min), n=10 mice. Two-sided Mann Whitney’s unpaired u test, Active: (st=0, ***p= 0.00074), Cylinder: (st= 22.5, ^ns^p= 0.1898). Two-sided Wilcoxon’s signed-rank paired test, Control: (st=1.0, *p= 0.045), Ablated: (st=0.0, **p= 0.0050). **M.** Individual mice representation of data in I for Control (blue lines) and Ptf1a neuron-ablated (red lines) animals. Data presented as mean ± SEM. See also Figure S3.

The level of activity, measured as speed of the animals during the circadian cycle, oscillated accordingly with the changes in lighting, increasing during the dark and decreasing during the light phases for all conditions (Figure 4B, 4G, S3H-I). We did not find significant changes between control and Ptf1a neuron-ablated animals’ average speed independently of the cycle phase (Figure S3G). Time spent scratching changed in agreement with the lighting phase for both control and Ptf1a-ablated animals, except at the end of the experiment where Ptf1a-ablated animals had developed peak scratching behavior (Figure 4C-F, S3E-F).

These findings led us to postulate a link between the level of activity of the animal and the intensity of the scratching phenotype since movement of the animal unescapably would lead to self-activation of its own hairs. A clear correlation between both variables was observed for control animals (Figure 4H) and to a lesser extent for Ptf1a neuron-ablated animals (Figure 4I), due to increased variability in scratching behavior compared to controls. We evaluated the scratching frequencies in two different contexts: as previously performed in their own home cages with the lid removed (as in Figure 3), which represents an active state given that the animal continuously explores the now open cage, and 30 minutes later in their own home cages once returned to the stabulation rack, which represents a resting state. Ptf1a neuron-ablated animals in the active context scratched 4-times more often than in the resting context (Figure 4J). Control animals also reduced scratching in the resting context, but at a much reduced level (Figure 4J). Each individual animal tested decreased its scratching frequency in the resting context (Figure 4K).

These results are in agreement with the behavior in the long-term analysis presented above. However, it was possible that higher scratching in the first (active) context was due to increased anxiety and not locomotion. To rule out this possibility, we placed mice in the same holding cylinder used for air puff delivery (as in Figure 3J), clearly a context with higher anxiety potential than the open home cage. Ptf1a neuron-ablated animals reduced scratching behavior more than 60-fold when placed in the cylinder, a context in which they can still move and scratch, but not locomote freely (Figure 4L-M). Taken together, these results argue in favor of the idea that spontaneous scratching observed after Ptf1a neuron ablation is an exacerbated response to self-stimulation of fur produced during normal activity.

### GRPR^+^ neurons elicit itch in Ptf1a neuron-ablated mice

To begin investigating if GRPR neurons are involved in the scratching phenotype of Ptf1a neuron-ablated animals, we inhibited GRPR neuron activity with a GRPR specific antagonist (Sukhtankar et al., 2013; Pereira et al., 2015) in Ptf1a neuron-ablated animals. Administration of the antagonist after removal of the bedding material from the home cage further reduced the scratching frequency of Ptf1a neuron-ablated animals by 50% (Figure 5A). In agreement with a model in which GRPR neuronal activity is involved in the scratching behavior of Ptf1a neuron-ablated mice, we found that 86% of vGlut1^+^ terminals contacting GRPR neurites received vGAT^+^ presynaptic inhibitory contacts most likely coming from Ptf1a neurons (Figure 5B). To obtain genetic evidence for such a role of GRPR neurons, we compared the scratching phenotype of Ptf1a neuron-ablated mice with mice in which both Ptf1a and GRPR neurons were ablated. Ablation efficiency of GRPR neurons was partial (Figure 5C-E), but sufficient to markedly decrease the scratching response to chemical pruritogens (Figure 5F, G). In agreement with previously reported data, GRPR neuron-ablated animals did not show any other sensory or motor deficiencies (Figure S4) (Sun and Chen, 2007; Sun et al., 2009). Interestingly, simultaneous ablation of Ptf1a and GRPR neurons delayed the development of the peak scratching activity by more than one day compared to Ptf1a neuron-ablated animals (Figure 5H), while having no effect on the peak scratching frequency (Figure 5I). Moreover, the increased sensitivity of Ptf1a neuron-ablated hairy skin towards gentle air puffs returned to normal in double-ablated animals (Figure 5J). In summary, these results suggest that Ptf1a neurons gate mechanosensory information in the GRPR pathway and that GRPR neurons elicit itch in response to this type of somatosensory input.

**Figure 5.**
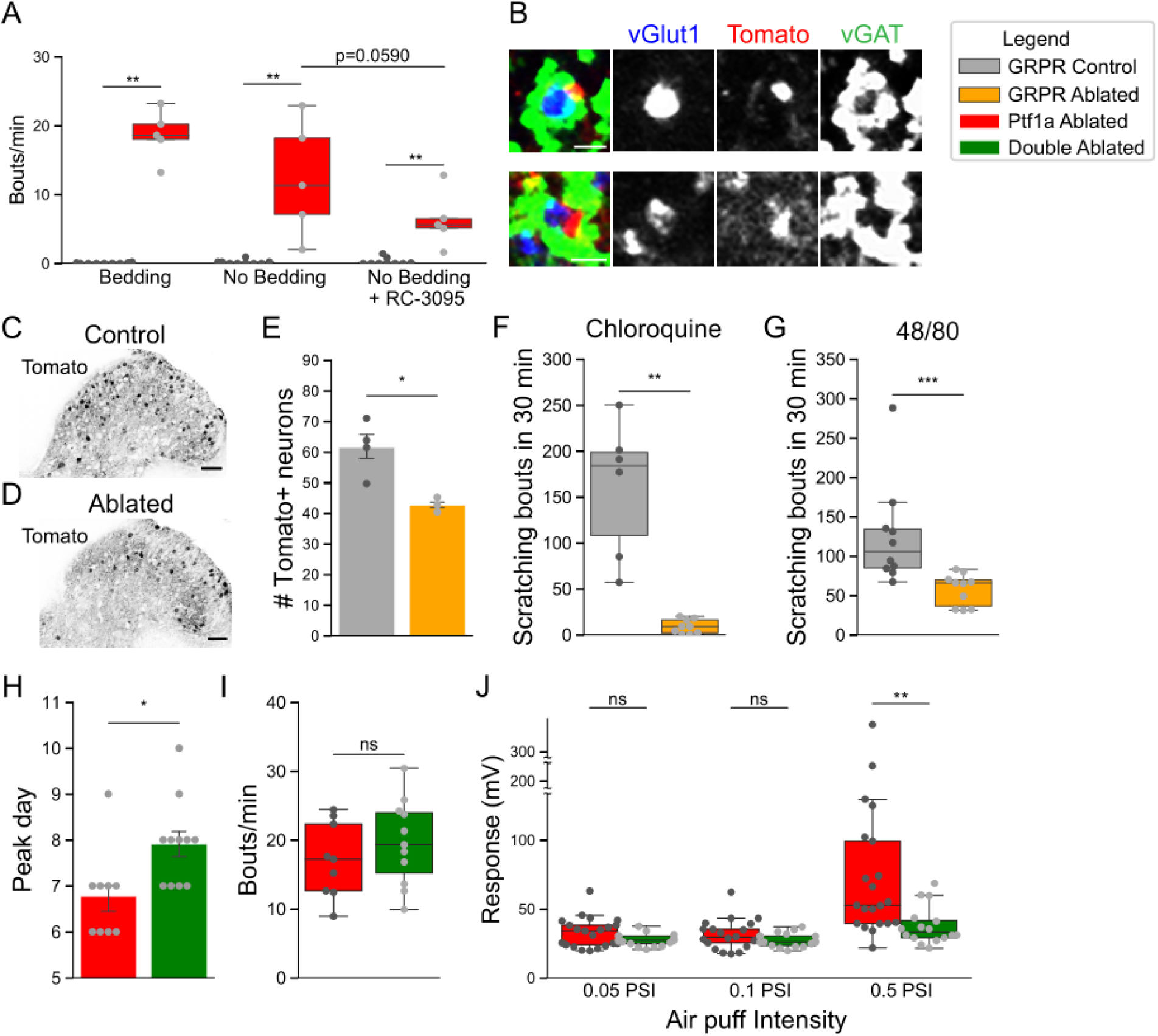
Spinal GRPR^+^ neurons mediate Ptf1a neuron-ablated scratching. **A.** GRPR antagonist RC-3095 effects on scratching frequency in control (blue) and Ptf1a neuron-ablated (red) animals. Bedding: Control (0.044±0.024 bouts/minute, n=9 mice), Ptf1a neuron-ablated (18.64±1.63 bouts/minute, n=5 mice); No bedding: Control (0.166±0.097 bouts/minute, n=9 mice), Ptf1a neuron-ablated (12.3±3.75 bouts/minute, n=5 mice); No bedding + RC-3095: Control (0.25±0.167 bouts/minute, n=9 mice), Ptf1a neuron-ablated (6.32±1.82 bouts/minute, n=5 mice). Two-sided Mann Whitney’s unpaired u test, Bedding: (st=0.0, **p=0.0022), No bedding: (st=0.0, **p=0.0026), No bedding + RC-2095: (st=0.0, **p=0.0023). Two-sided Wilcoxon’s signed-rank paired test, Ptf1a neuron-ablated No bedding vs Ptf1a neuron-ablated No bedding+RC-3095: (st=0.0, ^ns^p=0.0590). **B.** Representative images of GRPR-Cre; Ai9^lsl-tdTomato^ neurites contacting vGlut1^+^ somatosensory terminals apposed with vGAT^+^ inhibitory puncta. 86.6±5.2% of Tomato^+^ processes contacted by vGlut^+^ terminals show vGAT^+^ puncta (n=10-17 terminals/section, 3 sections/animal, 3 animals). **C, D.** Representative images of adult thoracic spinal cord hemisection showing GRPR neurons expressing Tomato before (**C**) and two weeks after (**D**) DT administration. **E.** Numbers of Tomato^+^ neurons in spinal dorsal hemisections in control (grey, 61.5±4.43 cells, n=4 mice) and GRPR neuron-ablated (orange, 42.7±1.4 cells, n=3 mice) animals. Two-sided Student’s unpaired t test, (st=3.495, *p=0.017). **F, G.** Quantification of scratching bouts in control (grey) and GRPR neuron-ablated (orange) animals after injection of chloroquine (**F**) or compound 48/80 (**G**) in the back. **F.** Control: 160.16±30.14 bouts, n=6 mice, Ablated: 8.66±2.6 bouts, n=9 mice. Two-sided Mann Whitney’s unpaired u test, (st=54.0, **p=0.0017). **G**. Control: 125±20.6 bouts, n=10 mice, Ablated: 57.5±6.33 bouts, n=10 mice. Two-sided Mann Whitney’s unpaired u test, (st= 94.5, ***p= 0.00087). **H.** Quantification of time after DT1 in which Ptf1a neuron-ablated (red) and Ptf1a;GRPR neuron-ablated (green) animals reached maximum scratching frequency. Ptf1a neuron-ablated: 6.7±0.32 days postDT, n=9 mice, Ptf1a;GRPR neuron-ablated: 7.9±0.28 days postDT, n=11 mice. Two-sided Mann Whitney’s unpaired u test, (st= 17.5, *p= 0.0122). **I.** Quantification of scratching frequency in Ptf1a neuron-ablated (red) and Ptf1a;GRPR neuron-ablated (green) animals in home cages, at peak day after DT treatment. Ptf1a neuron-ablated: 17.12±1.8 bouts/minute, n=9 mice, Ptf1a;GRPR neuron-ablated: 19.63±1.87 bouts/minute, n=11 mice. Two-sided Mann Whitney’s unpaired u test, (st=36.5, ^ns^p=0.34). **J.** Quantification of the activity response elicited by air puffs of increasing intensity directed towards the back of Ptf1a neuron-ablated (red, as in Figure 3D) and Ptf1a;GRPR neuron-ablated (green) animals 5 days after DT1. n = 17-21 animals per group. 0.05PSI: Ptf1a neuron-ablated 32.8±2.4 mV vs Ptf1a;GRPR neuron-ablated 27.4±1.06 mV; 0.1PSI: Ptf1a neuron-ablated 31.4±2.3 mV vs Ptf1a;GRPR neuron-ablated 27.4±1.22 mV; 0.5PSI: Ptf1a neuron-ablated 81.4±15.3 mV vs Ptf1a;GRPR neuron-ablated 37.7±3.4 mV. Two-sided Mann Whitney’s unpaired u test, ^ns^p>0.05, **p<0.01. Scale bar: B: 1 µm; Rest: 50 µm. Data presented as mean ± SEM See also Figure S4.

### Ptf1a neurons inhibit GRPR-mediated itch

To assess the function of Ptf1a neurons in controlling GRPR-mediated itch, we used a gain-of-function intersectional approach, where GRPR and/or Ptf1a neurons were activated with the activating DREADD receptor hM3Dq (Sciolino et al., 2016) (Figure 6A and S5A-B). As previously, we used littermates lacking one of the recombinase alleles treated with clozapine-N-oxide (CNO) as control animals. CNO-mediated activation of GRPR neurons induced a 20-fold increase in the number of scratching bouts compared to vehicle-treated or CNO-treated control animals (Figure 6B), as recently reported using a different approach (Bardoni et al., 2018). Chemogenetic activation of inhibitory Ptf1a neurons did not elicit any changes in somatosensory or motor behavior, nor in a conditioned place aversion assay (Figure S5C-K). Interestingly, simultaneous activation of Ptf1a and GRPR neurons, resulted in a 70% reduction in the number of scratching bouts compared to GRPR neuron-activated animals (Figure 6B). Together, these findings suggest that Ptf1a neurons gate mechanosensory information in the GRPR-mediated itch circuit.

**Figure 6.**
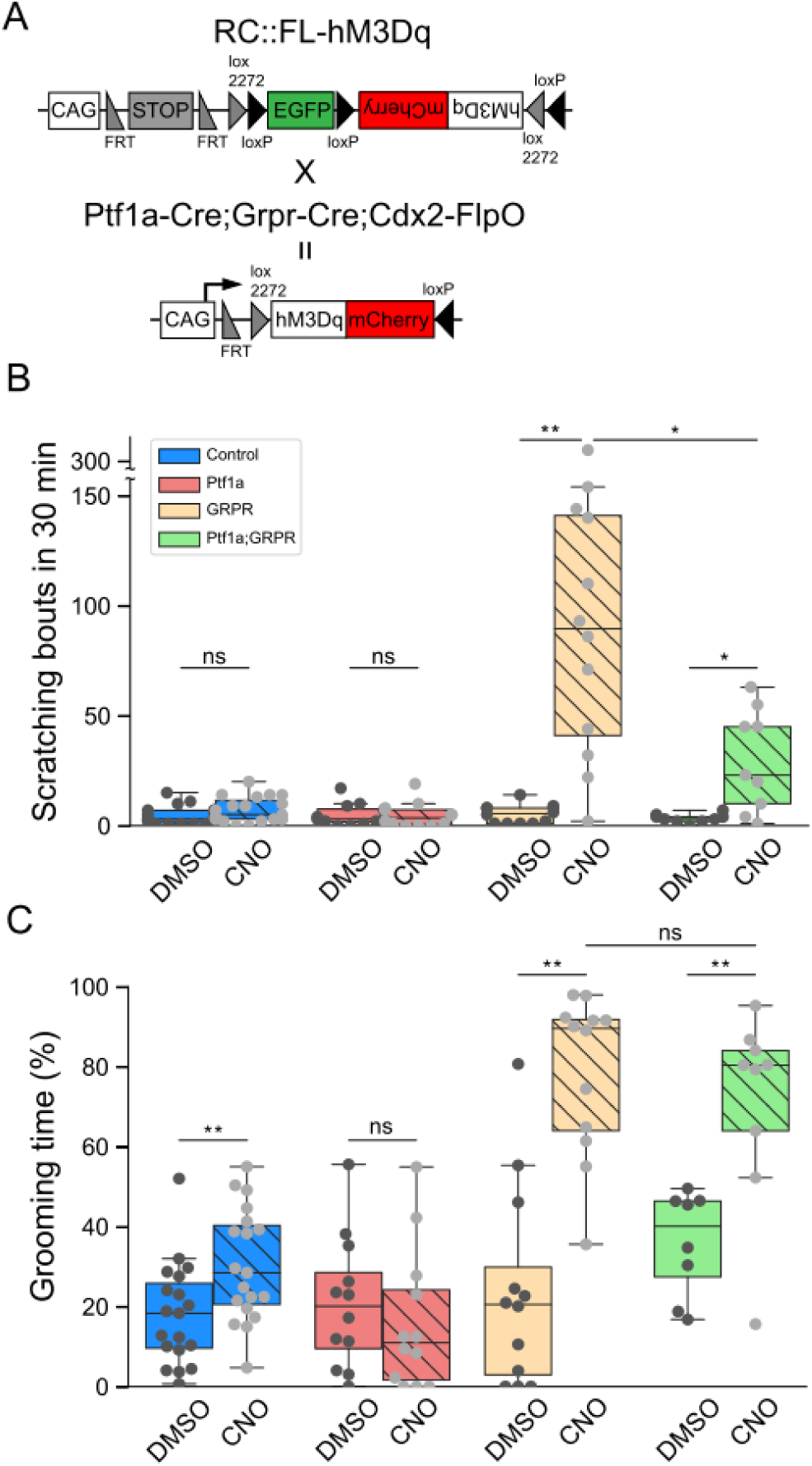
Ptf1a^+^ neuron activation inhibits GRPR neuron-induced itch. **A.** Schematic representation of genetic crosses for DREADD-mediated simultaneous activation of Ptf1a^+^ and GRPR^+^ neurons in the spinal cord. **B.** CNO-mediated activation of GRPR^+^ neurons induces scratching (DMSO: 5.33±1.22 bouts vs CNO: 100.25±23.53 bouts, n=12 mice; Two-sided Wilcoxon’s signed-rank paired test, st=1.0, **p=0.0033) which can be reduced through simultaneous activation of GRPR^+^ and Ptf1a^+^ neurons (DMSO: 3.33±0.67 bouts vs CNO: 29.55±7.66 bouts, n=9 mice; Two-sided Wilcoxon’s signed-rank paired test, st=1.0, *p=0.021; CNO: GRPR 100.25±23.53 bouts, n=12 mice vs Ptf1a;GRPR 29.55±7.66 bouts, n=9 mice, Two-sided Mann Whitney’s unpaired u test, st=87.0, *p=0.021). No differences are observed upon CNO administration to control (DMSO: 4.68±0.92 bouts vs CNO: 7.10±1.32 bouts, n=19 mice; Two-sided Wilcoxon’s signed-rank paired test, st=58.0, ^ns^p=0.24)) or Ptf1a neuron-activated animals (DMSO: 4.83±1.46 bouts vs CNO:4.83±1.63, n=12 mice; Two-sided Wilcoxon’s signed-rank paired test, st=33.0, ^ns^p=1.0) compared to the same animals treated with vehicle DMSO. **C.** Intense grooming induced by CNO-mediated activation of GRPR^+^ neurons (DMSO: 23.7±7.3% vs CNO: 78.5±5.8%, n=12 mice; Two-sided Wilcoxon’s signed-rank paired test, st=1.0, **p=0.0033) is not compensated for by the concomitant activation of Ptf1a^+^ neurons (DMSO: 36.1±4.6% vs CNO: 71±8.1%, n=9 mice; Two-sided Wilcoxon’s signed-rank paired test, st=2.0, *p=0.017; CNO: GRPR 78.5±5.8%, n=12 mice vs Ptf1a;GRPR 71±8.1%, n=9 mice, Two-sided Mann Whitney’s unpaired u test, st=69.0, ^ns^p=0.303). CNO-treated control animals spend a bit more time grooming (DMSO: 18±2.9% vs CNO: 30.4±3.2%, n=19 mice) which is suppressed by activation of Ptf1a^+^ neurons (DMSO: 20.8±4.7% vs CNO: 16.1±5.13%, n=12 mice; Two-sided Wilcoxon’s signed-rank paired test, st=28.0, ^ns^p=0.41). Data presented as mean ± SEM. See also Figure S5.

Grooming has recently been connected to a neuronal itch circuit (Gao et al., 2019). We observed that activation of GRPR neurons not only resulted in increased itch but also in increased grooming behavior. GRPR-activated animals spent 80% of the assay time grooming, compared to vehicle treatment (23%) (Figure 6C). Interestingly, simultaneous activation of GRPR and Ptf1a neurons did not result in a significant reduction of grooming activity, suggesting that those Ptf1a neurons that are targeted by Ptf1a-Cre do not control the GRPR-mediated grooming circuit. In summary, our results suggest a well-defined function for Ptf1a neurons in gating specific mechanosensory input to the GRPR itch circuit.

## Discussion

Chronic itch is a very common disease and secondary symptom originating from different causes (from dermatological to oncogenic) that remains without cure nowadays. Treatments are lacking and the burden for the patient and society is comparable to that of chronic pain (Kini et al., 2011). In this study, we report that spinal presynaptic inhibitory neurons marked by the expression of the transcription factor Ptf1a gate self-generated mechanosensory information during movement to prevent innocuous stimuli from triggering a scratch response (Figure 7). Loss of these neurons gives rise to a chronic itch phenotype and hairy skin hypersensitivity. We propose that presynaptic inhibition of cutaneous afferents in the spinal dorsal horn normally silences self-produced hair activation to facilitate adequate responses to external stimuli.

**Figure 7.**
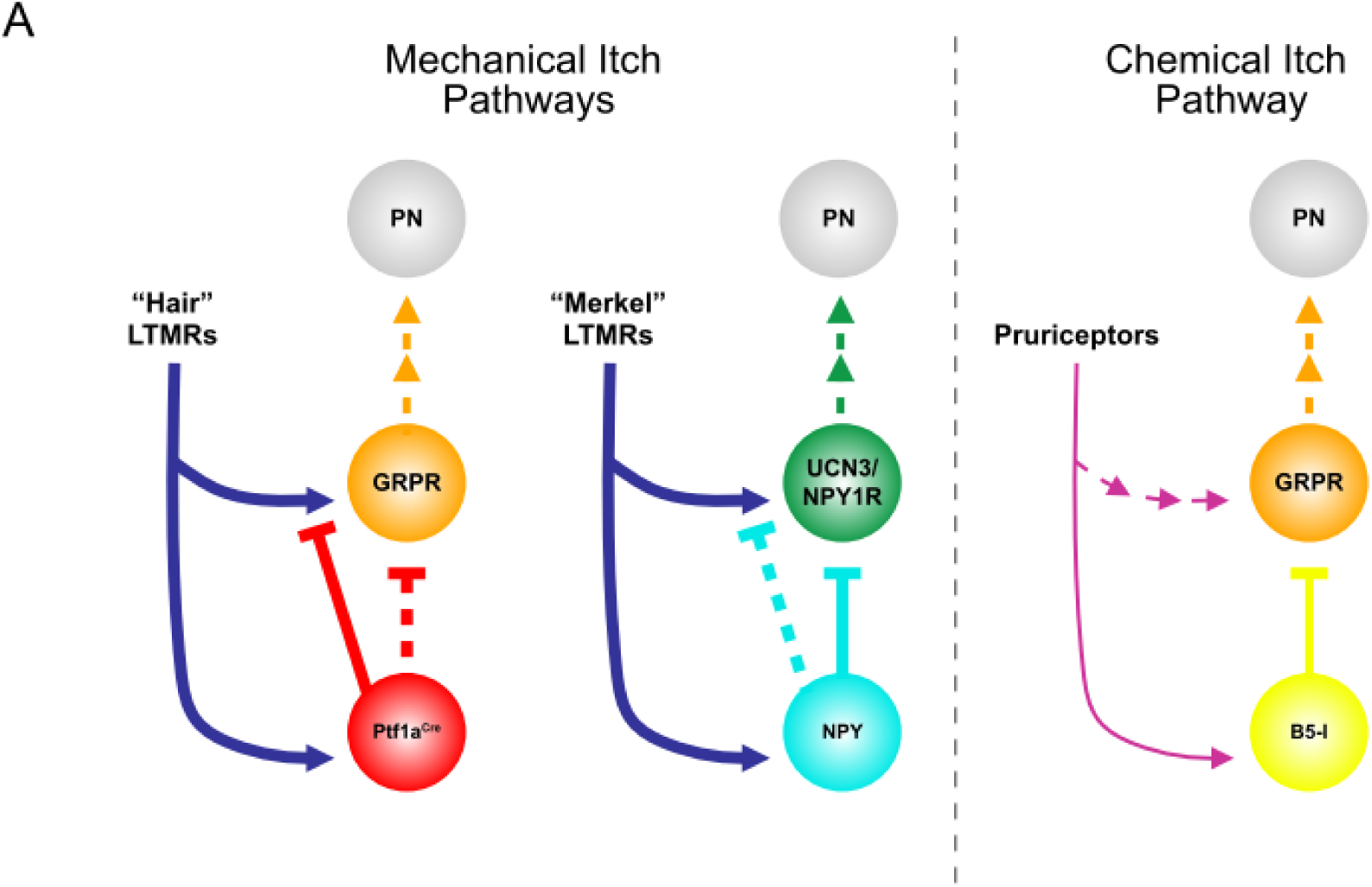
Itch models. Scheme representing a simplified view of itch circuits in the spinal dorsal horn according to the literature and the presented results. Inhibitory neurons that are lost in Bhlhb5 mutant mice (B5-I^+^) gate the chemical itch pathway exerting inhibition upon GRPR^+^ neurons, the central hub of chemical itch in the dorsal spinal cord. Mechanosensory information from low threshold mechanoreceptors (LTMRs) gate NPY^+^ neurons driving inhibition onto UCN3^+^ and NPY1R^+^ neurons. NPY+ contacts were previously described upon cutaneous mechanosensory afferents in the dorsal horn (Betley et al., 2009). Ptf1a-Cre neurons gate GRPR+ neuron activation through mechanosensory input driven by hair movement. The identity of LTMRs and whether they are shared between both mechanical pathways remains to be investigated (for a detailed molecular description of chemical vs mechanical itch pathways see Jakobsson et al., 2019). Projection neurons (PN) transmit itch signals to the brain. Serial arrows depict polysynaptic pathways. Dashed lines represent hypothetical connections based on the literature.

### Common developmental origin of spinal inhibitory neurons gating itch circuits

Our findings shed light on the complex specification of inhibitory neurons in the spinal cord. Ptf1a acts as a selector gene to promote a spinal inhibitory fate and Ptf1a knockout mice lack GABAergic neurons in the dorsal horns (Glasgow et al., 2005). However, Ptf1a knockout mice die perinatally (Glasgow et al., 2005; Hori et al., 2008) precluding any conclusions regarding neuronal subtype composition of the mature dorsal horns.

Previous work identified GAD65, GLYT2 and NPY as adult spinal inhibitory neurons subtypes whose ablation leads to the development of spontaneous scratching (Fink et al., 2014; Foster et al., 2015; Bourane et al., 2015a). Each of these genes have been shown to require Ptf1a for proper expression in the dorsal horns during spinal neurogenesis (Huang et al., 2008; Wildner et al., 2013; Borromeo et al., 2014). Such dependency establishes a developmental link for these subtypes to the dI4/dILA class of inhibitory neurons (Gross et al 2002, Müller et al 2002, Cheng et al 2004, Glasgow et al 2005, Wildner et al 2006). We reasoned that a common specification pathway during development might be relevant for their function in the mature dorsal horn circuits. Indeed, all three genes have been identified as markers for presynaptic inhibitory neurons: GAD65 is believed to be important for these neurons to sustain a high inhibitory tone (Hughes et al., 2005; Betley et al., 2009; Fink et al., 2014) and GLYT2 and NPY are markers for presynaptic inhibitory neurons targeting cutaneous afferents (Betley et al., 2009). Sharing such characteristics, we hypothesized that using Ptf1a as a genetic entry point to manipulate the function of these inhibitory neurons together would reveal the mechanisms of presynaptic inhibition control over the itch response.

Here we used a Ptf1a Cre-knockin mouse line, Ptf1a-Cre^EX1^ (Nakhai et al., 2007), with better recombination efficiency than the line most commonly used in spinal cord studies, Ptf1a-Cre^CVW^ (Kawaguchi et al., 2002). Our results show that Ptf1a-Cre^EX1^ targets ∼77% of inhibitory neurons expressing Pax2, whereas several of the well-known markers of spinal inhibitory neurons, including enkephalin, galanin or NPY, are not affected after Ptf1a-ablation. This is in agreement with recent single cell transcriptomic studies that identified at least six subtypes of dI4/dILA neurons in the developing spinal cord and only one of them was characterized by expression of Ptf1a (Delile et al., 2019). Consistently, NPY appeared as a marker of a different subtype (see also below).

As shown in our behavioral analysis, loss of Ptf1a neurons results in the most intense spontaneous scratching phenotype reported to date (Ptf1a ablation: 22 bouts/minute, GAD65 ablation: 5 bouts/minute, NPY ablation: 4.2 bouts/minute) and is also the fastest to develop after DT treatment (Ptf1a: 3.7 days, GAD65: 7 days (estimated), NPY: 1-2 weeks) (Fink et al., 2014; Bourane et al., 2015a; Acton et al., 2019). Our results suggest that Ptf1a captures a more complete set of the inhibitory subtypes involved in mechanical itch gating, resulting in a stronger phenotype. It also emphasizes the importance of presynaptic inhibitory neurons as an essential part of the cutaneous mechanosensory gating mechanisms that impinge onto spinal itch circuits.

### Presynaptic inhibition gates self-generated mechanosensory information

Our results showing that Ptf1a neurons make up for more than 80% of all presynaptic inhibitory contacts onto cutaneous mechanosensory fibers are in agreement with previous studies that characterized the developmental origin of spinal presynaptic inhibitory neurons (Betley et al., 2009) and their role in motor control gain (Fink et al., 2014). Presynaptic inhibition at the spinal cord level is a fundamental mechanism for the correct development of tactile sensitivity and adult social interactions among others (Orefice et al., 2016). A recent study explored spinal presynaptic inhibition in depth and elaborated on the importance of this form of sensory input control in hairy skin sensitivity (Zimmerman et al., 2019). Strikingly, the link between both areas of studies, namely Ptf1a neurons, the neurons responsible for such presynaptic inhibition at somatosensory afferents, have been so far missed. Here we show that loss of Ptf1a neurons increases hairy skin sensitivity which leads to the development of a dramatic spontaneous scratching phenotype including severe skin lesions. Our analysis of such phenotype in the home cages, which represent a naturalistic environment (Datta et al., 2019), suggests that loss of presynaptic inhibition of mechanosensory afferents underlies the increased hairy skin sensitivity that ultimately gives rise to scratching. The positive correlations of animal activity with scratching frequency plus that of hair stimulation with increased dorsal horn neural activity, suggest that self-generated activation of hair mechanosensory inputs during movement triggers an exacerbated scratch response in the absence of Ptf1a neuron inhibition. The presence of additional mechanosensory cues in the environment, like bedding material in the home cages, increases the likelihood of hair activation. In humans, clothes contact with skin hairs represents an equivalent scenario. Lack of an acute mechanical itch phenotype as the one previously described for NPY neuron-ablated animals (Bourane et al., 2015a; Acton et al., 2019; Pan et al., 2019) agrees with our Ptf1a-Cre targeting data, which reveals unaffected expression of NPY in the dorsal horns of Ptf1a neuron-ablated animals. Moreover, acute mechanical itch has been proposed to be mediated by Merkel touch-dome cells in the skin (Feng et al., 2018; Pan et al., 2019), which transmit a different type of mechanosensory information than hair mechanoreceptors. These results point to diverse subsets of presynaptic inhibitory neurons specialized in the gating of different mechanosensory information submodalities (Figure 7). Further studies are warranted to molecularly characterize the subtype of inhibitory neurons that is lost upon Ptf1a-Cre mediated ablation.

Thus, Ptf1a neurons seem to be part of a system in charge of blocking self-generated somatosensory stimuli during ongoing motor activity. Such a system has been shown to receive higher motor commands to reduce somatosensory gain during planning and execution of limb movements (Chapman and Tremblay, 2015; Juravle et al., 2017; Voudouris and Fiehler, 2017). Even though, throughout this work, we have used the term “spontaneous” scratching out of consistency with previous studies, we believe this scratching behavior should be redefined in order to avoid unnecessary confusion and given the existence of a stimulus to which the animal reacts, even if self-induced. We propose the use of “self-generated” scratching behavior, as a possible alternative.

### Itch and the perception of touch

Our discovery that DREADD-mediated activation of spinal GRPR neurons leads to increased scratching and grooming together with a delayed appearance of scratching phenotype in the simultaneous Ptf1a and GRPR neurons ablation paradigm, begs the question about the connection between itch and innocuous tactile information processing. Previous studies reported no changes in response to acute mechanical itch stimuli presented to GRPR neuron-ablated mice (Bourane et al., 2015a; Pan et al., 2019). However, such type of stimuli presentation, in which skin is pre-conditioned by removing hair before the experiment, would reduce hair mechanosensory input to the GRPR circuit, probably masking any outcome in response to hair activation.

Concomitant ablation of GRPR with Ptf1a neurons delayed the development of the scratching phenotype for more than one day. These data indicate that GRPR neurons normally receive direct or indirect mechanosensory cutaneous information. It has been recently shown that GRPR neurons require a strong excitatory drive from GRP neurons to elicit action potential firing (Pagani et al., 2019). Considering that 90% of body surface in mammals is covered by hairy skin (Djouhri et al., 2016), it is tempting to speculate that mechanosensory input to the circuit is used to maintain GRPR neurons at a high subthreshold level of excitation. Some evidence for this mechanism comes from the observation that GRPR neuronal activation (Figure 6B and Bardoni et al., 2018) evokes not only scratching behavior but also intense grooming activity (Figure 6C). It remains to be tested, albeit difficult to accomplish, the behavioral response elicited by GRPR neuronal activation in the absence of cutaneous mechanosensory information.

## Conclusion

The spinal circuit described here sheds light on how a subtype of inhibitory neurons gates self-generated mechanosensory information during ongoing motion. Malfunction of this mechanism evolves in a chronic itch syndrome reminiscent of many human pathologies where absence of perceived external stimulus ignites a strong itch-scratch cycle. Despite recent efforts toward the identification of the remaining itch circuit components, many essential questions remain to be solved in order to improve and design new therapies to fight such a debilitating disease as chronic itch.

## Acknowledgments

We thank the MPI animal facility for support in mouse line maintenance; Martyn Goulding (Salk Institute), Patricia Jensen (NIEHS), Silvia Arber (Biozentrum–University of Basel, Friedrich Miescher Institute), Jens Siveke (Technischen Universität München), Benjamin Arenkiel (Baylor College of Medicine) and Malin Lagerström (Uppsala Universitet) for mouse lines; Thomas Müller and Carmen Birchmeier (MDC Berlin) for antibodies; Kenneth Klau and Lisa Koletsou for assistance with animal facility work; Daniel del Toro, Hakan Kucukdereli and Amelia Douglass for constructive discussions and Sonia Paixão for critical reading of the manuscript. This study was supported by funds from the Max-Planck Society to RK and LaCaixa Banking Foundation through the Postdoctoral Junior Leader Fellowship Programme LCF/BQ/PI18/11630005 to AE.

## Author contributions

Conceptualization: A.E. and R.K.; Methodology: A.E.; Validation: A.E. and R.K.; Formal Analysis: A.E.; Investigation: A.E.; Resources: A.E.; Data Curation: A.E. and R.K.; Writing–Original Draft: A.E. and R.K.; Writing–Review and Editing: A.E. and R.K.; Visualization: A.E.; Supervision: R.K.; Funding Acquisition: A.E. and R.K.

## Materials and Methods

### Mouse lines

The mouse lines used in this study were previously described: Ptf1a-Cre^EX1^ (also known as p48^Cre^) (Nakhai et al., 2007); Ptf1a-Cre^CVW^ (Kawaguchi et al., 2002); Tau^lsl-SynaptophysinGFP^ (Tripodi et al., 2011); Ai9^lsl-tdTomato^ (B6.Cg-Gt(ROSA)26Sor^tm9(CAG-tdTomato)Hze^/J) (Madisen et al., 2010); Cdx2-FlpO (Britz et al., 2015); Lmx1b-Cre (Li et al., 2010); Tau^ds-DTR^ (Britz et al., 2015); Rosa26^CAG-ds-hM3Dq^ (Sciolino et al., 2016); Ai65D^ds-tdTomato^ (B6;129S-Gt(ROSA)26Sor^tm65.1(CAG-tdTomato)Hze^/J) (Madisen et al., 2015); RΦGT (Takatoh et al, 2013); GRPR-iCre (Aresh et al., 2017). Animals were kept and used in accordance with regulations from the government of Upper Bavaria. All lines were back-crossed to C57BL/6J mice at least 5 generations.

### Monosynaptic rabies virus tracings

Adult mice were anesthetized with isofluorane (2%) and placed in a stereotaxic frame (SR-5M-HT, Narishige) with a heating pad. Anesthesia was maintained throughout the procedures at 1.5% with a nose mask (GM-4, Narishige). The analgesics Metamizol (0.2 mg/g bodyweight, oral) and Carprofen (5mg/kg bodyweight, subcutaneous) were administrated. After incision of the skin and removal of dorsal intervertebral muscles, the vertebral column was fixed with a spinal column clamp (STS-A, Narishige). A laminectomy was performed at the C2 or L1 levels, the dura matter was carefully perforated with a fine needle to expose the spinal cord. 300 nl of virus were injected unilaterally on the dorsal spinal cord 500 μm lateral to the medial artery with a fine glass capillary using a microcontroller. The skin was then closed using a Reflex 7 skin closure system and tissue adhesive (3M Vetbond). Animals were perfused 6 days post-infection. The EnvA G-deleted rabies-GFP (Wickersham et al., 2006) were produced at the Salk Gene Transfer unit?.

### Histology

Animals were deeply anesthetized with ketamine/xylazin (100 mg/kg and 16 mg/kg respectively) and transcardially perfused first with phosphate-buffered saline (PBS) and then with 4% paraformaldehyde (PFA) in PBS. Brains and spinal cords were dissected, postfixed at 4°C in 4% PFA overnight, washed in PBS and embedded in 4% agarose. 70- to 100-μm sections were cut with a vibratome (Leica) and sections stored in cryoprotective solution (40% PBS, 30% glycerol, 30% ethylene glycol) at −20°C until use. Embryonic spinal cords were dissected and fixed in 4% PFA in PBS for 2h and processed as described above. DRGs were cryopreserved sequentially in 15% and 30% sucrose in PBS at 4°C, embedded in O.C.T (Fisher Scientific) and cut in a cryostat (Leica). 30 μm sections were mounted on slides, air dried and stored at −80°C.

### Immunohistochemistry

For immunofluorescence, vibratome free-floating sections and cryosections were blocked and permeabilized with 5% donkey serum, 0.3% TritonX-100 in PBS for 2 hours at room temperature (RT), following incubation with primary antibodies overnight at 4°C in 0.3% TritonX-100 in PBS. The following primary antibodies were used: goat anti-Pax2 (1:500) (AF3364, R&D Systems), guinea pig anti-Lmx1b (1:10,000) (gift from C. Birchmeier, Max Delbrück Center for Molecular Medicine, Germany), rabbit anti-cFos (1:2000) (sc-52, Santa Cruz), rabbit monoclonal anti-cFos (1:2000) (2250S, Cell signaling), rat anti-mCherry (1:1500) (M11217, Thermo), chicken anti-GFP (1:2000) (GFP-1010, Aves-Labs), chicken anti-βgalactosidase (ab9361, Abcam), anti-NF (1:1000) (NA1211, Affiniti), goat anti-TrkC (1:500) (AF1404, R&D Systems), sheep anti-TH (1:1000) (AB1542, Millipore), guinea pig anti-vGluT2 (1:1000) (AB2251, Millipore), guinea pig anti-vGluT1 (1:1500) (AB5905, Millipore), rabbit anti-DsRed (1:1000) (632496, Clontech), goat anti-CGRP (1:500) (36001, Abcam), rabbit anti-NPY (1:1000) (T-4070.0050, Peninsula Laboratories), goat anti-nNOS (1:1000) (ab1376, Abcam), rabbit anti-vGAT (1:1000) (131002, Synaptic Systems), rabbit anti-ENK (1:1000) (20065, Immunostar), mouse anti-GAD65 (1:50) (GAD6, Developmental Studies Hybridoma Bank, created by the NICHD of the NIH and maintained at The University of Iowa, Department of Biology, Iowa City, IA 52242), rabbit anti-Galanin (1:1000) (T-4334, BMA Biomed), rabbit anti-RFP (1:1000) (600-401-379, Rockland). After three 10-min washes in PBS, sections were incubated with secondary antibodies for 2 h at room temperature in 0.3% TritonX-100 PBS. The following secondary antibodies or counterstains were used: donkey anti-rabbit/mouse/goat/sheep/rat/chicken/guinea pig conjugated to Alexa Fluor 488, Cy3, Alexa Fluor 647 or Cy5 (1:500), Neurotrace 640/660 (1:200) (N-21483, Molecular Probes) or DAPI (1:1000). After three 15-min washes in PBS, sections were mounted in slides and coverslipped with mowiol mounting medium.

### Microscopy

Fluorescence z-stack images were acquired with an Olympus FV1000 or Leica SP8 confocal microscope. Images were minimally processed with FiJi/ImageJ software (NIH) or Adobe Photoshop to enhance brightness and contrast. No filters were used to decrease noise. Co-localization analysis quantification was done in single confocal z-sections. Measurements were performed using FiJi/ImageJ software.

For analysis of GFP co-localization with vGAT in presynaptic puncta we followed previously described methods (Abraira et al., 2017). In short, 20-60 dorsal horn randomly selected vGlut1/vGAT appositions were marked in ImageJ per hemisection without visualizing the GFP channel. Percentages of vGlut1/vGAT appositions in which the vGAT terminal was also GFP were calculated after revealing the GFP channel. For analysis of vGlut1/vGAT appositions in DT treated animals, 50 randomly selected vGlut1 terminals were marked in ImageJ per hemisection without visualizing the vGAT channel. Percentages of vGlut1 terminals apposed by vGAT synapses were calculated after revealing the vGAT channel.

### Spatial distribution of neurons

The position of interneurons in the spinal cord was recorded manually in FiJi/ImageJ. Cartesian coordinates for each interneuron were determined in the transverse spinal cord plane with respect to the midpoint of the central canal, defined as position (0,0). Custom Python scripts were used to calculate the probability density function and display two-dimensional kernel density estimations for each cell population. Estimates were graphically displayed as shaded or line contour plots, with lines connecting points of equal probability density.

### Pharmacological treatments

Cell ablation was induced by intraperitoneal administration of diphtheria toxin at 50ng/g of body weight (D0564, Sigma-Aldrich) in saline solution. Injection was performed twice with a 72 hours interval (Duan et al., 2014; Bourane et al., 2015a; Acton et al., 2019; Pan et al., 2019). GRPR antagonist RC-3095 (Tocris) was dissolved in 0.1% DMSO in saline to 2µg/µl and injected intraperitoneally at 10µg/g of body weight 30 minutes before behavioral evaluation. DREADD activation was induced by intraperitoneal injection of CNO (C0832, Sigma-Aldrich) at 2ng/g of body weight diluted in vehicle (2% DMSO in saline), behavioral tests were performed 30 minutes later.

### Behavioral assays

Animals were acclimated to the behavioral room and testing apparatus during three consecutive days for at least 30 minutes unless otherwise indicated. Efforts were made to remain blind to the genotype of the tested animals, however given the obvious nature of the phenotype in Ptf1a-ablated animals it was not always possible. Littermates of either sex were used in all tests. Except for the hot plate and rotarod, all the testing apparatuses were custom designed and built from acrylic. Drawings are available upon request.

#### Home cage video recording (as in Figure 4C, 4D, 4I, …)

Animals were singly housed at least one week before DT treatment. Before DT1, animals in their home cages were video recorded with cage lid and food tray removed. A transparent lamina of acrylic was used to prevent animals from getting out. Mice were recorded in pairs for 10 minutes. To evaluate the influence of bedding material in spontaneous scratching behavior, a clean cage without bedding was stacked inside the home cage to preserve as much as possible olfactory cues. The double cage was returned to the rack for 30 minutes and another video recording of 10 minutes was acquired. To evaluate the role of GRPR neurons in Ptf1a neuron-mediated scratching, mice were injected with the GRPR antagonist RC-3095, returned to the double cage and rack for 30 minutes and another video recording of 10 minutes was acquired. Both scratching frequency and grooming time were calculated offline. To evaluate the influence of animal motion on scratching behavior (Figure 5J-K), Ptf1a neuron-ablated mice at peak scratching frequency and control littermates were recorded in their home cages 30 minutes after returning them to the cage rack.

#### Chemical itch

Animals were lightly anesthetized with isofluorane and injected in the back skin at the level of thoracic vertebrae 12. Compound 48/80 (100 ug, Millipore) or chloroquine (200 ug, Millipore) in 50 uL of sterile saline were injected. Animals were video recorded for 30 minutes and the scratching bouts after injection counted.

#### Hairy skin sensitivity

To evaluate changes in hairy skin sensitivity at different air puff intensities we adapted a previously published system (Orefice et al., 2016). We used a SR-LAB startle response system (SDI-2325-0400, San Diego Instruments) adapted to deliver air puffs (SDI-2325-0368) to the back of the animal. Pressure regulation was achieved through a forward pressure reducing regulator (4136ANNKE, Equilibar). Animals were habituated for 10 minutes to the holding cylinder, noise and light levels the day before testing. Continuous white noise level was set to 90dB to prevent mice from hearing the opening of the valve. The protocol consisted of 5 min acclimation, followed by 7 trains of 5 air puffs each delivered at pseudorandom interstimulus intervals at increasing pressures from 0.05 PSI to 2.5 PSI.

#### cFos induction through hair stimulation

To assess the ability of hair movement to induce neuronal activity in the spinal dorsal horn, we adapted a previous protocol designed to stimulate the skin (Duan et al., 2014). We kept mice under light anesthesia (1-1.5% isofluorane) using a head mask. Manipulation of the animal was reduced to the minimum possible to avoid confounding stimulations. The back hairs of the animal were combed with a soft paintbrush (#6) rostrocaudally and caudorostrally from the forelimbs to the hindlimbs for 150 times in 5 minutes. This protocol was applied 3 times with 1 minute intertrials (450 strokes in total during 17 minutes). After stimulation, animals were kept under anesthesia for 2 hours, transcardially perfused and processed as previously described for cFos immunostaining.

#### Dynamic brush

Light touch sensitivity was evaluated with the dynamic light brush test (Bourane et al., 2015a). Mice were placed in individual compartments of a testing apparatus with a wire mesh floor. The plantar surface of the hindpaw was lightly stimulated with a soft brush (#6) stroking from heel to toe. Stimulation was repeated 10 times on alternating sides and the percentage of paw withdrawal were calculated.

#### Von Frey

This test was used to assess sensitivity to static touch and mechanical pain. Mice were placed in individual compartments of a testing apparatus with a wire mesh floor. Calibrated von Frey monofilaments (0.02-2 g) (Aesthesio Precision Tactile Sensory Evaluator) were used to stimulate the medial plantar surface of the hindpaws. Withdrawal, licking or biting following or immediately after the 5 seconds stimulus were considered as a positive response. Paw withdrawal threshold (PWT) were calculated according to (Detloff et al., 2012).

#### Pinprick

This test was used to evaluate mechanical pain. Mice were placed in individual compartments of a testing apparatus with a wire mesh floor. The plantar surface of the hindpaw stimulated with an Austerlitz insect pin (0.02 mm; Fine Science Tools). Stimulation was repeated 10 times on alternating sides and the percentage of paw withdrawal were calculated.

#### Dynamic hot plate

Heat sensitivity was evaluated through determination of the thermal withdrawal threshold. Mice were acclimated to the testing apparatus (IITC Life Science) set at 32°C the day before and the testing day for 10 minutes. The assay consisted on 3 trials in which the plate temperature increased from 32°C to 55°C at a rate of 3°C/minute. The ramp was stopped when the animals licked, shooked the hindpaw or jumped.

#### Cold plantar assay

Cold sensitivity was measured as previously described (Brenner et al., 2012). Mice were placed in individual compartments of a testing apparatus with a 4 mm glass floor. A pellet of crushed dry ice was positioned in contact with the glass beneath the hindpaw. The stimulus was presented 6 times on alternating sides. Latency to withdrawal was measured.

#### Acute mechanical itch

We performed two versions of this test, one applying the stimulus to the shaved nape of the neck and another one to the shaved back of the animal according to Bourane et al., 2015a. Mice were placed in individual compartments of a testing apparatus with a wire mesh floor and after 30 minutes habituation, received five separate stimuli for ∼1 second at random sites on the skin of the target region with a von Frey monofilament of 0.07g. Scratching of the poked site was considered a positive response.

#### Rotarod

An accelerating rotarod (Ugo Basile) test was used to evaluate gross motor coordination. Mice were trained on 2 consecutive days at a constant speed of 5 rpm for 5 minutes. The training was repeated until each animal stayed in the cylinder without falling down for the whole 5 minutes. The third day the rotarod was programmed to accelerate from 5 to 40 rpm in 5 minutes. Each animal was subject to 3 trials with an intertrial recovery of 15 minutes. Latency to fall, maximum speed and distance walked were registered for each animal.

#### Open field

Overall locomotor activity and anxiety was evaluated in a custom-made acrylic 40×40×25 cm square arena. Mice were free to explore it for 15 minutes. Mean speed, distance traveled and time spent in the center of the arena (defined as a central 20×20 cm area) was measured with Ethovision XT 11 (Noldus).

#### Startle reflex/Prepulse inhibition (PPI)

PPI testing was performed essentially as a previously described (Orefice et al., 2016; Orefice et al., 2019) with some modifications. The tests were carried out on the same setup used for testing hairy skin sensitivity. Here we evaluated the ability of a tactile pre-stimulus (0.9 PSI, 50 ms) to inhibit startle to a stronger tactile stimulus (3 PSI, 5 ms) (tactile-tactile PPI test) or to a loud acoustic stimulus (120 dB, 20 ms) (tactile-acoustic PPI test) with an interstimulus interval of 250 ms. We sought to investigate whether Ptf1a- or GRPR-ablated animals exhibit specific tactile sensorimotor gating deficits. Animals were habituated for 10 minutes to the holding cylinder, noise and light levels the day before testing. Both PPI test versions were carried out on different days. On testing day, mice were placed inside the ventilated, cylindrical holder on a platform within a soundproof chamber. Protocols were as described (Orefice et al., 2016): acclimation phase with constant white noise (90 dB) for 5 minutes, block I, block II, block III and block IV trials. All stimuli were pseudorandomly spaced between 15 and 45 seconds. Block I consisted of 5 startle stimuli alone (3 PSI 5 ms or 120 dB white noise for tactile-tactile and tactile-acoustic sessions, respectively), to measure the initial startle reflex. Block II consisted of 5 prepulse stimuli alone (0.9 PSI 50 ms), to measure response to the prepulse stimulus alone. Block III incorporated prepulse/pulse, pulse alone and no stimulation trials. Block IV consisted of 5 startle stimuli alone, to measure habituation. Whole body flinch, or startle reflex, was quantified using an accelerometer sensor measuring the amplitude of movement of the animal within the cylindrical holder.

#### Long-term surveillance of mice in home cages

The plans for the modified cage lid, infrared LED array, camera, lens, raspberry pi and acquisition code are available upon request. We modified the cage lid for adapting a NoIR PiCamera with a modified M12 220° lens (RP-VC1N, Entaniya) to have direct view of the whole inner space of the home cage. The camera was attached to a Raspberry Pi 3 Model B (Raspberry Pi Foundation) running Raspbian and the picamera python interface was used to control acquisition and saving of the video stream. An acrylic enclosure was custom built to fit around the home cage and fitted with 5 meters of an 850 nm infrared LED strip (Solarox IR1-60-850, LED1.de). Acrylic diffusers were set in place between the LED strip and the cage walls. Mice were habituated for 24 hours to the enclosure after DT2 and recording started. Video recording was stopped when Ptf1a-animals reached peak scratching frequency. FFmpeg was used to convert raw h264 movie streams to compressed video. A neural network was trained on 2000 frames labelled manually with the position and orientation of the mouse to generate an accurate elipse-fit around the mouse (Geuther et al., 2019) and track all the videos. Output of the trained neural network was used to generate the input data necessary for JAABA (Kabra et al., 2013) and train a behavior classifier to identify frames in which mice engaged in scratching behavior with the hindpaws. The resulting behavior classifier had an accuracy of 88.4% for scratch frames and 78% for non-scratch frames in videos it had never been trained on (ground-truth). This classifier was applied to all the videos for each animal. Scratching time and speed of the animals were computed (Figure S3A-B).

#### Scratching in a locomotion-restrictive context (Figure 5L-M)

To evaluate the influence of animal movement on scratching frequency, Ptf1a neuron-ablated mice at peak scratching frequency and control littermates were recorded in their home cages (see above) and 30 minutes later inside the same holding cylinder used in the startle reflex system. Animals were habituated for 10 minutes the day before testing and recorded for 10 minutes on testing day. Scratching frequency was quantified offline.

#### Conditioned Place Aversion (CPA)

To test for the valence associated with chemogenetic activation of Ptf1a-derived neurons we performed a CPA test as described in (Vrontou et al., 2013). We designed a custom made acrylic 2-chamber arena (each chamber 15×15×30 cm, connected through a 5×5 cm opening that could be closed). Each chamber was designed with a different pattern of alternating black and white stripes (vertical or horizontal) and different patterned floor to help mice to distinguish each chamber during conditioning. On the first day, mice were free to explore both chambers for 30 minutes (Pre-Test). Conditioning started on the second day, confining the mouse to the initially preferred (IP) chamber with CNO treatment for 30 minutes. The third day, the mouse was confined to the initially non-preferred chamber (INP) after administration of vehicle. The fourth and fifth day, the cycle was repeated. On the sixth day, the mouse was allowed to explore freely both chambers and preference was calculated (Post-Test). Percentage of time spent in each chamber was assessed using Ethovision XT 11 software (Noldus).

#### Chemogenetic induction of itch and grooming

To evaluate itch and grooming induced behaviors by DREADD-mediated activation, mice treated with vehicle or CNO in consecutive days were placed in individual compartments of a testing apparatus with a wire mesh floor. A bottom-up view of the animals was video recorded for offline quantification. Scratching bouts and grooming time were calculated for 30 and 10 minutes, respectively, 30 minutes after treatment.

### Statistical analysis

No statistical methods were used to predetermine sample sizes. Data presented as bar graphs indicate mean ± SEM (standard error of the mean). Dots in bar graphs and box plots represent individual values per mouse, line indicates median. Statistical analyses were performed in Python and MATLAB using custom scripts. Significance levels are indicated as follows: ^ns^p> 0.05; *p< 0.05; **p< 0.01; ***p< 0.001; ****p< 0.0001.

---

**Figure S1.**
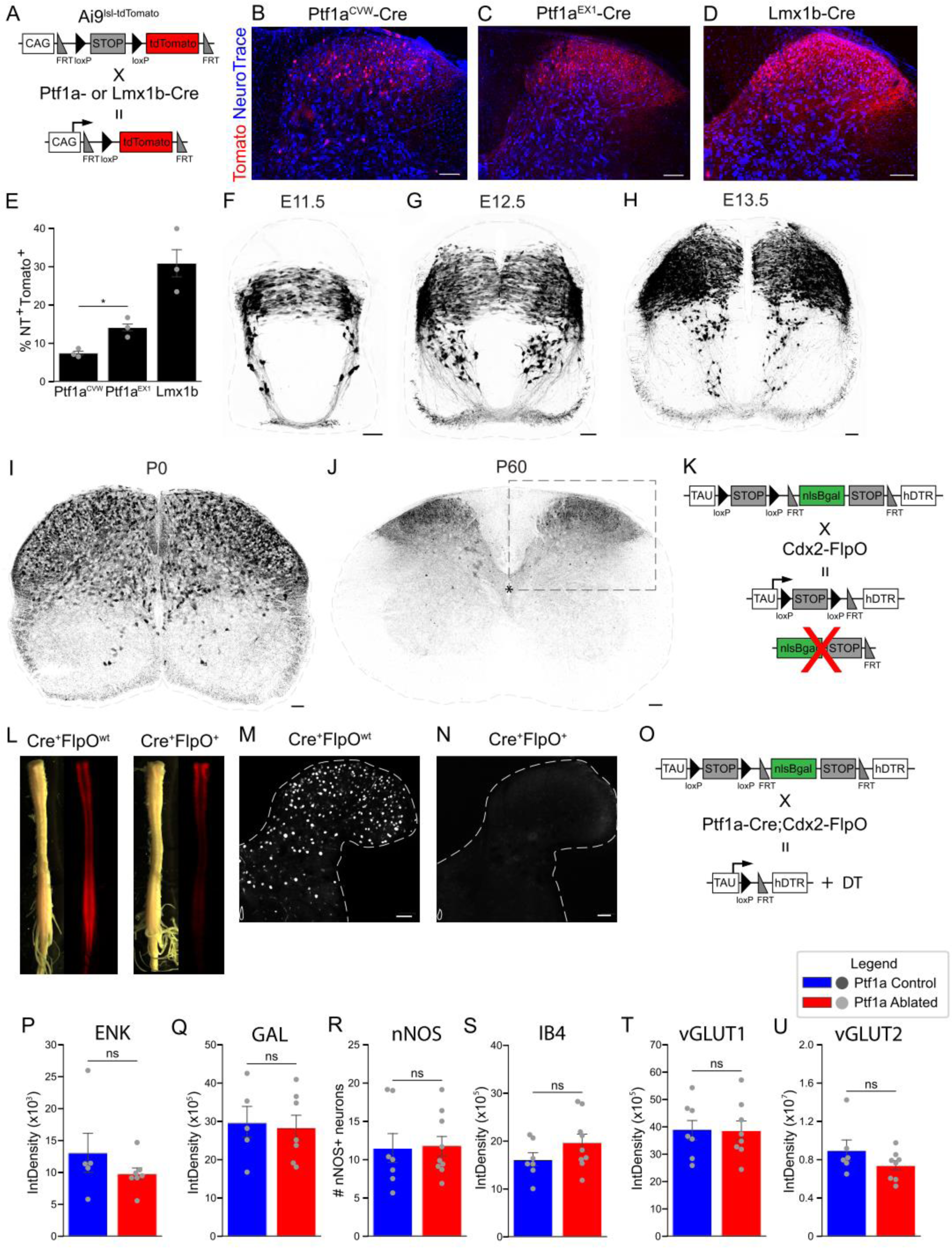
Characterization of Ptf1a-Cre mouse lines, ontogenesis of adult dorsal horn neurons and Ptf1a neurons as GABApre adult neurons (related to Figure 1). **A.** Scheme representing the genetic strategy used for lineage tracing of Ptf1a- and Lmx1b-derived neurons. **B-D.** Dorsal horn region of adult Ptf1a-Cre^CVW^; Ai9^lsl-tdTomato^ (**B**), Ptf1a-Cre^EX1^; Ai9^lsl-tdTomato^ (**C**) and Lmx1b-Cre; Ai9^lsl-tdTomato^ (**D**), respectively, used for quantification. Immunofluorescence for Tomato and NeuroTrace (NT) as neuronal marker. **E.** Quantification of percentage of NT^+^ neurons in adult spinal sections derived from Ptf1a-Cre^CVW^; Ai9^lsl-tdTomato^ (7.38±0.64%), Ptf1a-Cre^EX1^; Ai9^lsl-tdTomato^ (14±1.5%) and Lmx1b-Cre; Ai9^lsl-tdTomato^ (31±5%) mice. Excitatory dorsal horn neurons labelled with Lmx1b-Cre (Li et al., 2010; Alaynick et al. 2011) were 2-fold more abundant than Ptf1a-Cre^EX1^ neurons using the same reporter mouse line; n=3 hemisections/animal, 3 animals/genotype. Two-sided Student’s unpaired t test, (st=−4.18, *p=0.014). **F-J.** Developmental course of Ptf1a neurons in Ptf1a-Cre^EX1^; Ai9^lsl-tdTomato^ mice. At early stages, neurons originating from the mid-dorsal progenitor domain migrate mediolaterally towards the lateral early alar plates. These cells constitute the dI4/dILA class of dorsal spinal neurons (Alaynick et al., 2011). A subset of early Tomato^+^ neurons migrate ventrally towards the floor plate. At adult stages, no Tomato^+^ neurons are found ventral to the central canal (asterisk). Inset shows area used for quantification in E. **K.** Scheme representing strategy used to direct selective expression of hDTR in the spinal cord. Cells expressing FlpO recombinase lose expression of βgal, which allows assessment of Cre and FlpO co-expression in the same cell. **L.** Spinal cords from Ptf1a-Cre;Cdx2-FlpO;Ai9^lsltdTomato^ animals positive for Cre (left panel) or both Cre and FlpO (right panel), together with Ai9^lsltdTomato^. Cdx2-FlpO mouse line drives expression caudal to C5 as it can be observed by the loss of Tomato fluorescence, due to the removal of the FRT-flanked Ai9 cassette (Madisen et al., 2010) (in each panel: left, visible light; right, tomato fluorescence). **M, N.** Representative images of βgal immunofluorescence in spinal cord sections from Ptf1a-Cre;Cdx2-FlpO;Tau^ds-DTR^ animals positive for Cre (**M**) or both Cre and FlpO (**N**). Loss of βgal in Ptf1a neurons in the presence of Cdx2-FlpO demonstrates the efficiency of the system to specifically target all Ptf1a-Cre neurons in the spinal cord. **O.** Scheme representing strategy used for intersectional genetic DT-mediated ablation of spinal Ptf1a-Cre derived neurons. **P.** Quantification of Enkephalin immunofluorescence in the dorsal horn of control (1.3×10^4^±3.4×10^3^ AU) and Ptf1a neuron-ablated (9.7×10^3^±1.02×10^3^ AU) mice. n=10-12 hemisections/animal, 5-7 animals per group. Two-sided Student’s unpaired t test, (st= 0.927, ^ns^p= 0.3988). **Q.** Quantification of Galanin immunofluorescence in the dorsal horn of control (2.95×10^6^±4.4×10^5^ AU) and Ptf1a neuron-ablated neurons (2.82×10^6^±3.4×10^5^ AU). n=10-12 hemisections/animal, 7-9 animals per group. Two-sided Student’s unpaired t test, (st= 0.2287, ^ns^p= 0.8237). **R.** Quantification of the number of nNOS^+^ neurons in dorsal hemisections of the spinal cord in control (11.46±2.05 cells) and Ptf1a neuron-ablated (11.78±1.37 cells) animals. n=10-12 hemisections/animal, 7-9 animals per group. Two-sided Student’s unpaired t test, (st= −0.1349, ^ns^p= 0.8946). **S.** Quantification of IB4 immunofluorescence in the dorsal horn of control (1.6×10^6^±1.47×10^5^ AU) and Ptf1a neuron-ablated (1.96×10^6^±1.87×10^5^ AU) animals. n=10-12 hemisections/animal, 7-9 animals per group. Two-sided Student’s unpaired t test, (st= −1.4297, ^ns^p= 0.1747). **T.** Quantification of vGlut1 immunofluorescence in the dorsal horn of control (3.89×10^6^±3.91×10^5^ AU) and Ptf1a neuron-ablated (3.85×10^6^±3.65×10^5^ AU) animals. n=10-12 hemisections/animal, 6-8 animals per group. Two-sided Student’s unpaired t test, (st= 0.084, ^ns^p=0.934). **U.** Quantification of vGlut2 immunofluorescence in the dorsal horn of control (8.93×10^6^±1.1×10^6^ AU) and Ptf1a neuron-ablated (7.4×10^6^±5.4×10^5^ AU) animals. n=10-12 hemisections/animal, 6-8 animals per group. Two-sided Student’s unpaired t test, (st=1.27, p=0.24). Scale bars: B-D, J: 100 µm; F-I, M, N: 50 µm. Data presented as mean ± SEM.

**Figure S2.**
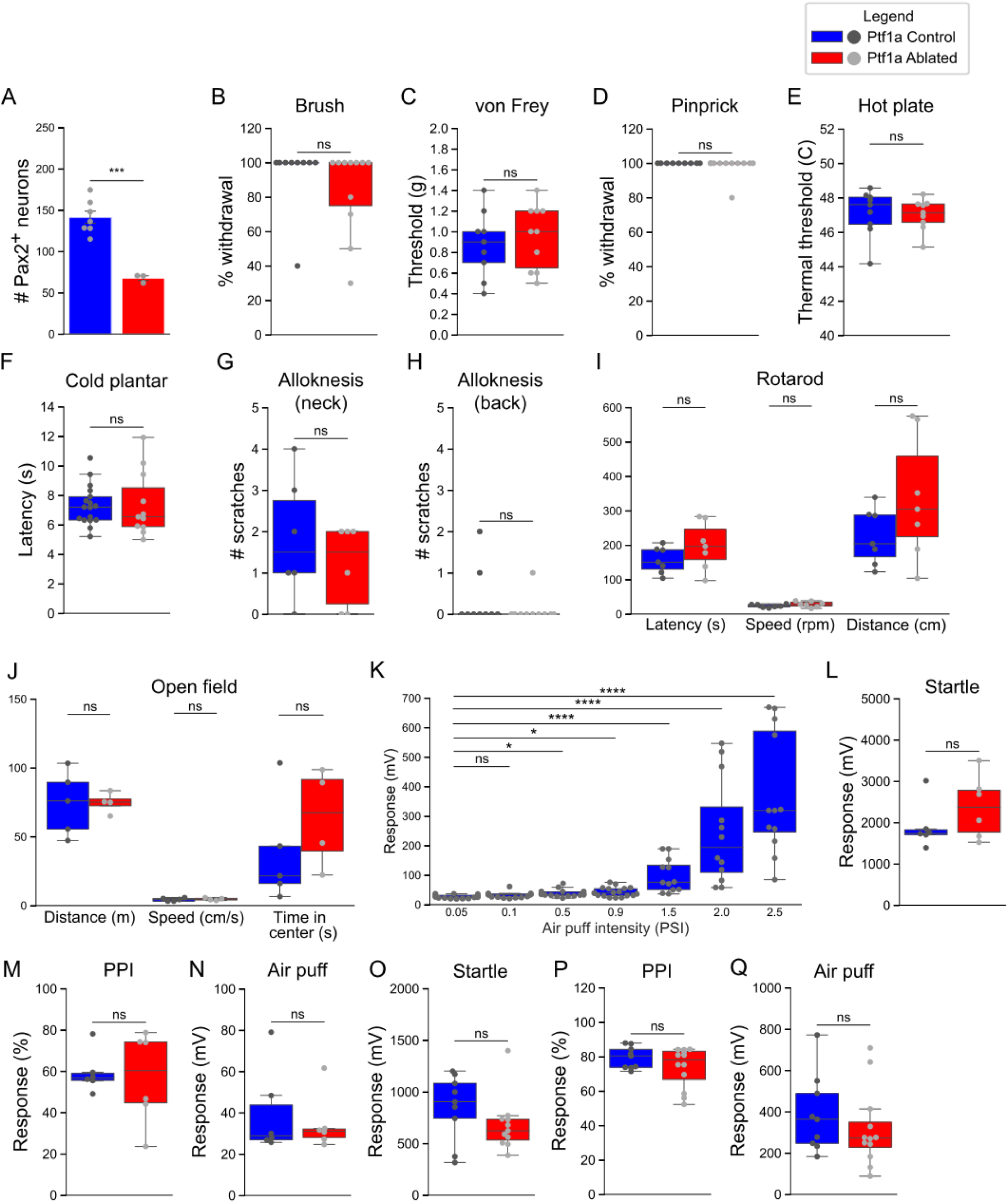
Ptf1a neuron-ablated animals show no defects in other sensory modalities (related to Figure 3). **A.** Quantification of the number of Pax2^+^ spinal neurons in dorsal hemisections of control (141.36±7.75 cells) and Ptf1a neuron-ablated (67.75±2.87 cells) animals 5 days after DT1. n= 3-5 hemisections/animal, 3-7 animals per group. Two-sided Student’s unpaired t test, (st=5.95, ***p=0.0003). **B.** No differences in dynamic brush test between control (blue, 93.33±6.66%, n=9 mice) and Ptf1a neuron-ablated (red, 84.54±7.43%, n=11 mice) animals 5 days after DT1. Two-sided Mann Whitney’s unpaired u test, (st= 61.0, ^ns^p= 0.272). **C.** No differences in von Frey paw withdrawal threshold between control (blue, 0.87±0.1, n=9 mice) and Ptf1a neuron-ablated (red, 0.95±0.1, n=10 mice) animals 5 days after DT1. Two-sided Student’s unpaired t test, (st= −0.499, ^ns^p= 0.623). **D.** No differences in pinprick test between control (blue, 100±0%, n=9 mice) and Ptf1a neuron-ablated (red, 98.18±1.81%, n=11 mice) animals 5 days after DT1. Two-sided Mann Whitney’s unpaired u test, (st= 54.0, ^ns^p= 0.421). **E.** No differences in hot plate test between control (blue, 47.13±0.45°C, n=9 mice) and Ptf1a neuron-ablated (red, 47.01±0.30°C, n=9 mice) animals 5 days after DT1. Two-sided Student’s unpaired t test, (st=0.216, ^ns^p=0.831). **F.** No differences in cold plantar test between control (blue, 7.32±0.33s, n=17 mice) and Ptf1a neuron-ablated (red, 7.36±0.66s, n=11 mice) animals 5 days after DT1. Two-sided Student’s unpaired t test, (st= −0.07, ^ns^p= 0.944). **G.** No differences in alloknesis test in shaved neck between control (blue, 1.83±0.6 scratches, n=6 mice) and Ptf1a neuron-ablated (red, 1.16±0.4 scratches, n=6 mice) animals 5 days after DT1. Two-sided Mann Whitney’s unpaired u test, (st= 22.5, ^ns^p= 0.508). **H.** No differences in alloknesis test in back between control (blue, 0.33±0.23 scratches, n=9 mice) and Ptf1a neuron-ablated (red, 0.1±0.09 scratches, n=10 mice) animals 5 days after DT1. Two-sided Mann Whitney’s unpaired u test, (st= 51.0, ^ns^p= 0.44). **I.** No differences in rotarod test between control (blue, latency: 156.47±14.38s, speed: 23.66±1.69 rpm, distance: 224.71±30.68 cm, n=7 mice) and Ptf1a neuron-ablated (red, latency: 197.71±25.96s, speed: 28.24±3.06 rpm, distance: 335.55±67.65 cm, n=7 mice) animals 5 days after DT1. Two-sided Student’s unpaired t test, Latency: (st= −1.389, ^ns^p= 0.1899), Speed: (st=−1.305, ^ns^p=0.216), Distance: (st=−1.49, p=0.161). **J.** No differences in open field test between control (blue, distance: 74.32±10.4 m, speed: 4.3±0.52 cm/s, time in center: 38.18±17.43s, n=5 mice) and Ptf1a neuron-ablated (red, distance: 74.65±3.77 m, speed: 4.64±0.45 cm/s, time in center: 63.92±18.1s, n=4 mice) animals 5 days after DT1. Two-sided Student’s unpaired t test, Distance: (st= −0.027, ^ns^p= 0.979), Speed: (st= −0.482, ^ns^p= 0.645), Time in center: (st= −1.015, ^ns^p= 0.344). **K.** Evaluation of the response of control animals to increasing intensities of air puffs directed to the back hairy skin: 0.05 PSI: 27.25±1.46 mV, 0.1 PSI: 32.41±2.2 mV, 0.5 PSI: 37.7±2.8 mV, 0.9 PSI: 42.03±3.73 mV, 1.5 PSI: 98.52±16.22 mV, 2.0 PSI: 243.83±51.31 mV, 2.5 PSI: 372.8±59.53 mV. n=12-18 animals per intensity. One-way Kruskal-Wallis test followed by post hoc Fisher’s least significance difference comparison: ^ns^p>0.05, *p<0.05, ****p<0.0001. **L-N.** No differences in tactile-acoustic startle response test between control (blue, startle: 1905.83±230.21 mV, PPI: 59.25±4.02%, prepulse: 39.6±8.6 mV, n=6 mice) and Ptf1a neuron-ablated (red, startle: 2374±309.5 mV, PPI: 56.88±9%, prepulse: 34.67±5.5 mV, n=6 mice) animals 5 days after DT1. Two-sided Mann Whitney’s unpaired u test, Startle: (st= 13.0, ^ns^p= 0.4711), Prepulse: (st=17.0, ^ns^p=0.936), PPI: (st=20.0, ^ns^p=0.81). **O-Q.** No differences in tactile-tactile startle response test between control (blue, startle: 849±107.25 mV, PPI: 79.4±2.11%, prepulse: 387.64±62.56 mV, n=9 mice) and Ptf1a neuron-ablated (red, startle: 679.3±80.2 mV, PPI: 73.67±3.42%, prepulse: 317.23±54.25 mV, n=12 mice) animals 5 days after DT1. Two-sided Mann Whitney’s unpaired u test, Startle: (st= 69.0, ^ns^p= 0.148), Prepulse: (st=68.0, ^ns^p=0.34), PPI: (st= 65.0, ^ns^p= 0.455). Data presented as mean ± SEM.

**Figure S3.**
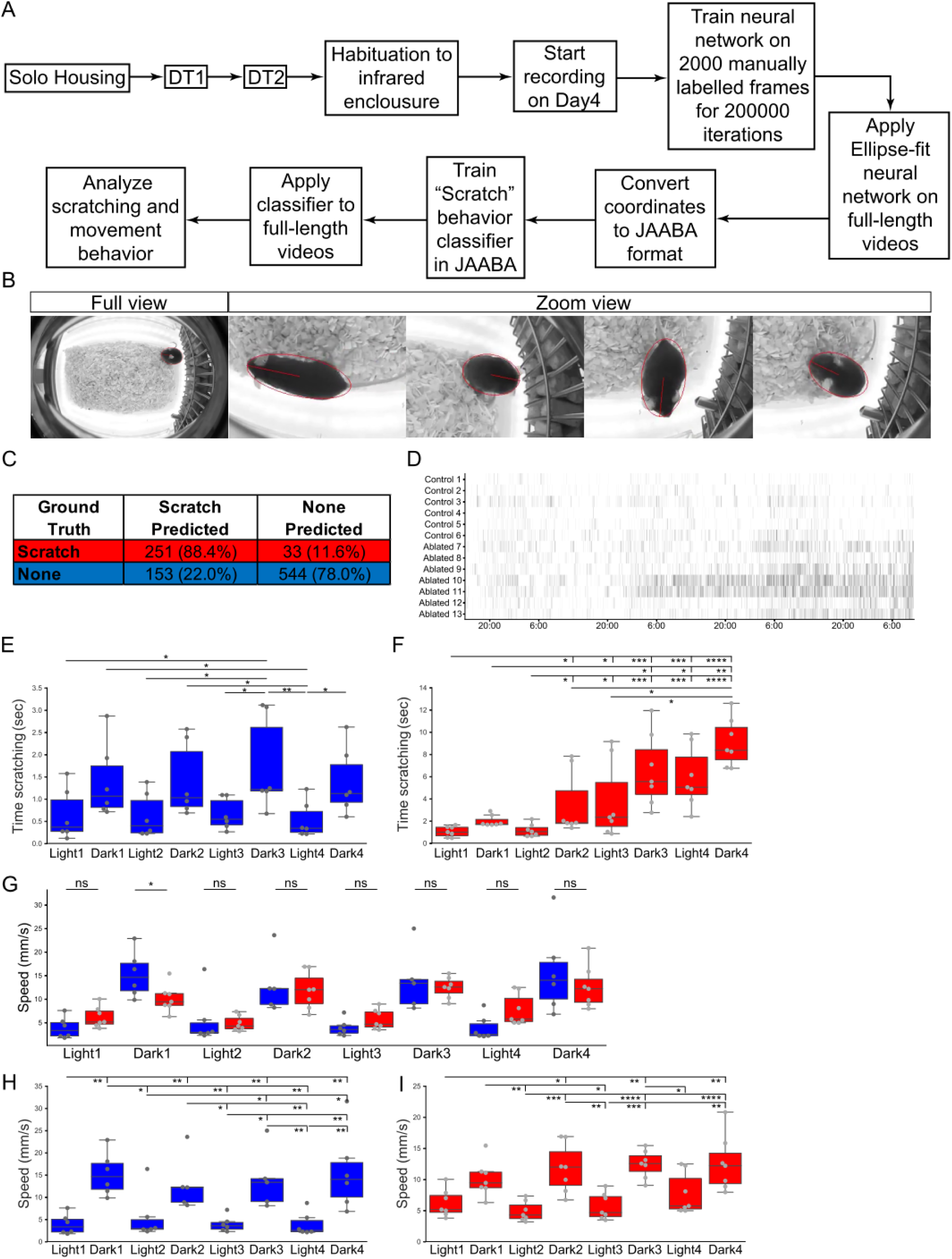
Overview and analysis of long-term naturalistic mouse scratching behavior (related to Figure 4). **A.** Protocol pipeline for acquisition, analysis and quantification of 90 hour data from mice in their home cages. **B.** Representative images of the outcome from the neural network trained to identify, segment and fit an ellipse to mice. Zoom view images depict instances of angles correctly assigned in the four directions. **C.** JAABA classification scores in ground-truth mode. **D.** Raster plot representation of individual scratch events for each control (n=6 mice) and Ptf1a neuron-ablated (n=7 mice) animals during the whole ∼90 hours of acquired video. **E, F.** Quantification of scratching time in control (**E**, 6 mice) and Ptf1a neuron-ablated (**F**, 7 mice) animals consecutive light/dark phases. One-way Kruskal-Wallis test followed by post hoc Fisher’s least significance difference comparison: *p<0.05, **p<0.01, ***p<0.001, ****p<0.0001. Note: Y axis scale in D different from E. **G.** Quantification of speed in different light/dark phases for control (blue, n=6 mice) and Ptf1a neuron-ablated (red, n=7mice) animals. No statistically significant differences: ns. Two-sided Mann Whitney’s unpaired u test: *p<0.05. **H, I.** Quantification of speed in control animals (**H**, 6 mice) and Ptf1a neuron-ablated animals (**I**, 7 mice) during consecutive light/dark phases. One-way Kruskal-Wallis test followed by post hoc Fisher’s least significance difference comparison: *p<0.05, **p<0.01, ***p<0.001, ****p<0.0001. Note: Y axis scale in I different from H.

**Figure S4.**
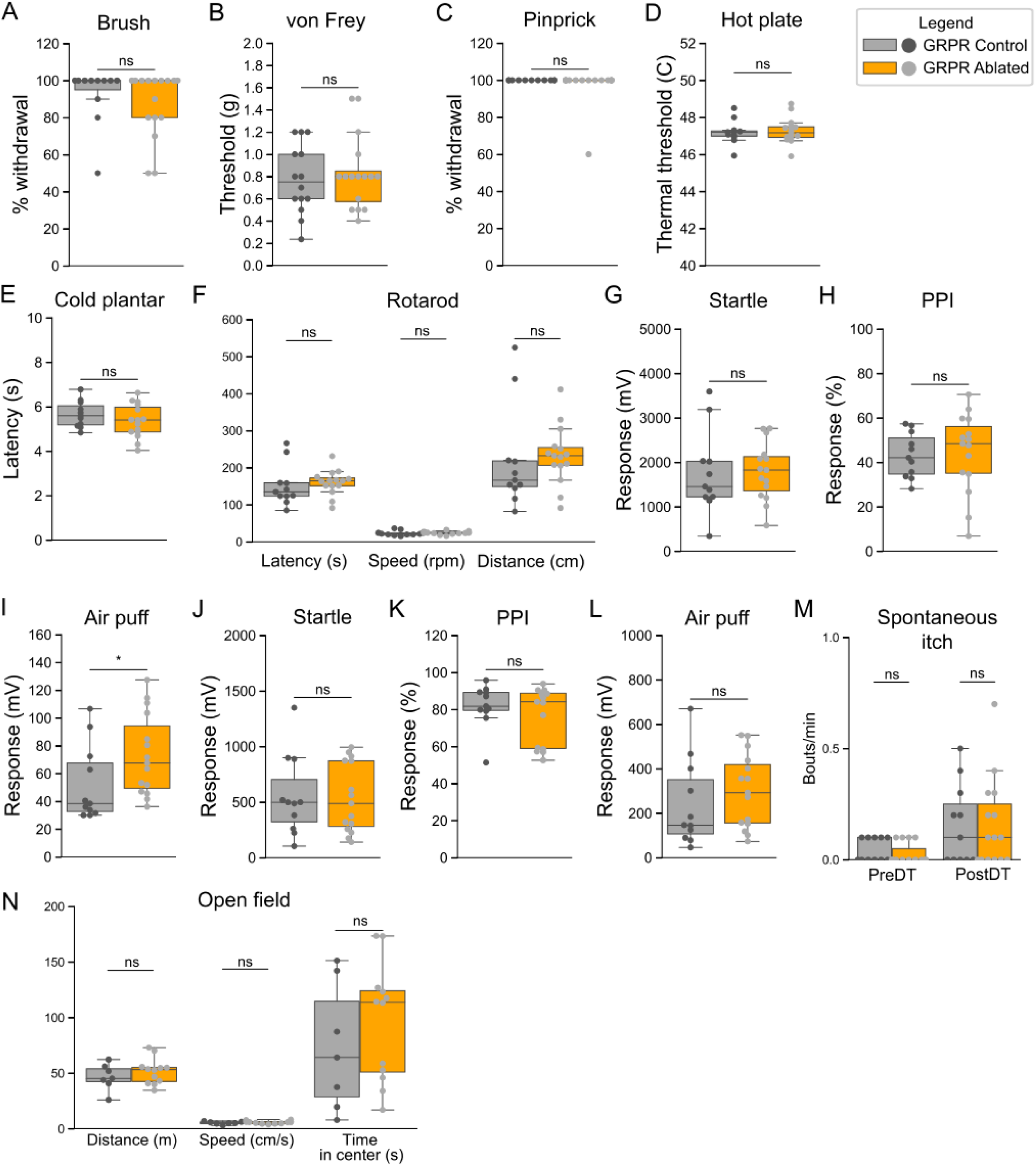
GRPR neuron-ablated animals show no defects in other sensory modalities (related to Figure 5). **A.** No differences in dynamic brush test between control (grey, 92.72±4.7 %, n=11 mice) and GRPR neuron-ablated (orange, 87.5±4.42 %, n=16 mice) animals 2 weeks after DT1. Two-sided Mann Whitney’s unpaired u test, (st=104.0, ^ns^p=0.376). **B.** No differences in von Frey paw withdrawal threshold between control (grey, 0.77±0.08 g, n=14 mice) and GRPR neuron-ablated (orange, 0.83±0.08 g, n=16 mice) animals 2 weeks after DT1. Two-sided Student’s unpaired t test, (st=−0.488, ^ns^p=0.63). **C.** No differences in pinprick test between control (grey, 100±0 %, n=11 mice) and GRPR neuron-ablated (orange, 97.5±2.5 %, n=16 mice) animals 2 weeks after DT1. Two-sided Mann Whitney’s unpaired u test, (st=93.5, ^ns^p=0.451). **D.** No differences in hot plate test between control (grey, 47.2±0.19 °C, n=11 mice) and GRPR neuron-ablated (orange, 47.27±0.17 °C, n=16 mice) animals 2 weeks after DT1. Two-sided Student’s unpaired t test, (st=−0.27, ^ns^p=0.78). **E.** No differences in cold plantar test between control (grey, 5.66±0.18 seconds, n=11 mice) and GRPR neuron-ablated (orange, 5.35±0.19 seconds, n=15 mice) animals 2 weeks after DT1. Two-sided Student’s unpaired t test, (st=1.11, ^ns^p=0.277). **F.** No differences in rotarod test between control (grey, latency: 151.3±16.7 seconds, speed: 22.88±1.97 rpm, distance: 218.37±41.48 cm, n=11 mice) and GRPR neuron-ablated (orange, latency: 159.97±8.54 seconds, speed: 24.06±1.04 rpm, distance: 232.66±20.32 cm, n=15 mice) animals 2 weeks after DT1. Two-sided Student’s unpaired t test, Latency: (st=−0.50, ^ns^p=0.62), Speed: (st=−0.57, ^ns^p=0.57), Distance: (st=−0.33, ^ns^p=0.74). **G-I.** No differences in tactile-acoustic startle response test between control (grey, startle: 1756.52±281.93 mV, PPI: 43.13±3.04%, prepulse: 52.33±8.27 mV, n= mice) and GRPR neuron-ablated (orange, startle: 1794.9±166.63 mV, PPI: 44.36±4.6%, prepulse: 73.3±7.55 mV, n= mice) animals 2 weeks after DT1. Two-sided Mann Whitney’s unpaired u test, Startle: (st=71.0, ^ns^p=0.57), Prepulse: (st=40.0, *p=0.029), PPI: (st=70.0, ^ns^p=0.53). **J-L.** No differences in tactile-tactile startle response test between control (grey, startle: 554.54±108.8 mV, PPI: 81.8±3.55 %, prepulse: 240.78±59.31 mV, n=11 mice) and GRPR neuron-ablated (orange, startle: 536.7±79.97 mV, PPI: 77.04±3.93 %, prepulse: 299.29±42.76 mV, n=15 mice) animals 2 weeks after DT1. Two-sided Mann Whitney’s unpaired u test, Startle: (st=83.0, ^ns^p=1.0), Prepulse: (st=62.0, ^ns^p=0.3), PPI: (st=90.0, ^ns^p=0.716). **M.** Scratching frequency in control (grey) and GRPR neuron-ablated (orange) animals in home cages, before and after DT treatment. Control (PreDT: 0.045±0.016 bouts/minute, n=11 mice, PostDT: 0.154±0.054 bouts/minute, n=11 mice), Ablated (PreDT: 0.026±0.012 bouts/minute, n=15 mice, PostDT: 0.16±0.05 bouts/minute, n=15 mice). Two-sided Mann Whitney’s unpaired u test, PreDT (st=98.0, ^ns^p=0.345), PostDT (st=82.0, ^ns^p=1.0). **N.** No differences in open field test between control (grey, distance: 46.52±4.45 m, speed: 5.17±0.5 cm/s, time in center: 72.75±21.57 s, n=7 mice) and GRPR neuron-ablated (orange, distance: 51.6±3.35 m, speed: 5.74±0.37 cm/s, time in center: 95.8±15.23 s, n=12 mice) animals after DT. Two-sided Student’s unpaired t test, Distance: (st=−0.917, ^ns^p=0.37), Speed: (st=−0.918, ^ns^p=0.37), Time in center: (st=−1.015, ^ns^p=0.34). Data presented as mean ± SEM.

**Figure S5.**
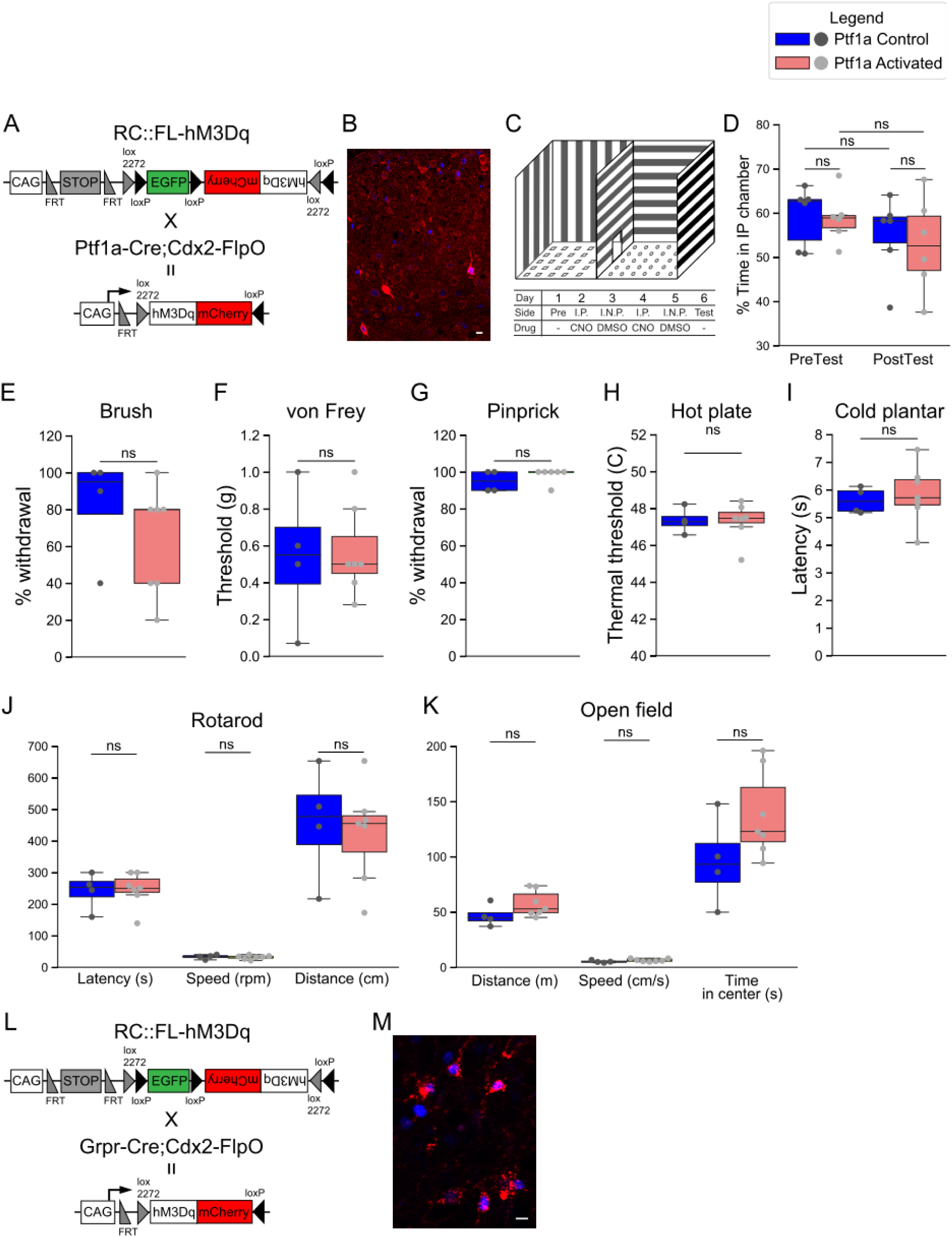
Ptf1a^+^ neuron-activation does not affect other sensory modalities (related to Figure 6). **A.** Schematic representation of genetic crosses for DREADD-mediated Ptf1a^+^ neuron activation in the spinal cord. **B.** Representative image of mCherry (red) and cFos (blue) immunofluorescence in Ptf1a neuron-activated animals after CNO treatment. **C.** Schematic representation of CPA assay apparatus and experimental timeline. **D.** CNO-mediated activation of Ptf1a^+^ neurons does not have an effect on conditioned place preference. Control (blue, PreTest: 59.4±2.73%, PostTest: 55.03±3.66%, n=6 mice), Excited (pink, PreTest: 58.8±2.3%, PostTest: 52.8±4.36%, n=6 mice). Two-sided Student’s unpaired t test, PreTest: (st=0.16, ^ns^p=0.87), PostTest: (st=0.38, ^ns^p=0.70). Two-sided Student’s paired t test, Control: (st=1.39, ^ns^p=0.22), Excited: (st=2.31, ^ns^p=0.069). **E.** No differences in dynamic brush test between control (blue, 82.5±14.36%, n=4 mice) and Ptf1a neuron-activated (pink, 62.8±11.06%, n=7 mice) animals 30 minutes after CNO administration. Two-sided Mann Whitney’s unpaired u test, (st=21.0, ^ns^p=0.21). **F.** No differences in von Frey paw withdrawal threshold between control (blue, 0.54±0.19 g, n=4 mice) and Ptf1a neuron-activated (pink, 0.57±0.09, n=17 mice) animals 30 minutes after CNO administration. Two-sided Student’s unpaired t test, (st=−0.14, ^ns^p=0.89). **G.** No differences in pinprick test between control (blue, 95±2.3%, n=4 mice) and Ptf1a neuron-activated (pink, 98.3±1.67%, n=6 mice) animals 30 minutes after CNO administration. Two-sided Mann Whitney’s unpaired u test, (st=8.0, ^ns^p=0.35). **H.** No differences in hot plate test between control (blue, 47.33±0.34°C, n=4 mice) and Ptf1a neuron-activated (pink, 47.3±0.39°C, n=7 mice) animals 30 minutes after CNO administration. Two-sided Student’s unpaired t test, (st=0.076, ^ns^p=0.94). **I.** No differences in cold plantar test between control (blue, 5.62±0.24s, n=4 mice) and Ptf1a neuron-activated (pink, 5.84±0.39s, n=7 mice) animals 30 minutes after CNO administration. Two-sided Student’s unpaired t test, (st=−0.40, ^ns^p=0.69). **J.** No differences in rotarod test between control (blue, latency: 241.46±29.7 s, speed: 33.12±3.48 rpm, distance: 456.08±90.83 cm, n=4 mice) and Ptf1a neuron-activated (pink, latency: 246.1±20.5 s, speed: 31.67±2.32 rpm, distance: 424.35±58.5 cm, n=6 mice) animals 30 minutes after CNO administration. Two-sided Student’s unpaired t test, Latency: (st=−0.13, ^ns^p=0.89), Speed: (st=0.36, ^ns^p=0.72), Distance: (st=0.31, ^ns^p=0.76). **K.** No differences in open field test between control (blue, distance: 46.68±4.93 m, speed: 5.19±0.55 cm/s, time in center: 96.01±20.26 s, n=4 mice) and Ptf1a neuron-activated (pink, distance: 57.56±4.4 m, speed: 6.4±0.49 cm/s, time in center: 137.95±14.77 s, n=7 mice) animals 30 minutes after CNO administration. Two-sided Student’s unpaired t test, Distance: (st=−1.566, ^ns^p=0.152), Speed: (st=−1.566, ^ns^p=0.152), Time in center: (st=−1.7, ^ns^p=0.125). **L.** Schematic representation of genetic crosses for DREADD-mediated GRPR+ neuron activation in the spinal cord. **M.** Representative image of mCherry (red) and cFos (blue) immunofluorescence in GRPR^+^ neuron-activated animals after CNO treatment. **N.** Scale bars: 10 µm. Data presented as mean ± SEM.

## References

Abraira, V.E., Kuehn, E.D., Chirila, A.M., Springel, M.W., Toliver, A.A., Zimmerman, A.L., Orefice, L.L., Boyle, K.A., Bai, L., Song, B.J., et al. (2017). The Cellular and Synaptic Architecture of the Mechanosensory Dorsal Horn. Cell 168, 295–310.e19.

Acton, D., Ren, X., Di Costanzo, S., Dalet, A., Bourane, S., Bertocchi, I., Eva, C., and Goulding, M. (2019). Spinal Neuropeptide Y1 Receptor-Expressing Neurons Form an Essential Excitatory Pathway for Mechanical Itch. Cell Reports 28, 625–639.e6.

Alaynick, W.A., Jessell, T.M., and Pfaff, S.L. (2011). SnapShot: Spinal Cord Development. Cell 146, 178–178.e1.

Aresh, B., Freitag, F.B., Perry, S., Blümel, E., Lau, J., Franck, M.C.M., and Lagerström, M.C. (2017). Spinal cord interneurons expressing the gastrin-releasing peptide receptor convey itch through VGLUT2-mediated signaling: PAIN 158, 945–961.

Azim, E., Fink, A.J.P., and Jessell, T.M. (2014). Internal and External Feedback Circuits for Skilled Forelimb Movement. Cold Spring Harbor Symposia on Quantitative Biology 79, 81–92.

Babayan, BM. and Conen, CS. (Eds.). (2019). Behavior matters [Special issue]. Neuron, 104, 1, p1–176.

Bardoni, R., Takazawa, T., Tong, C.-K., Choudhury, P., Scherrer, G., and MacDermott, A.B. (2013). Pre- and postsynaptic inhibitory control in the spinal cord dorsal horn: Inhibitory control in the dorsal horn. Annals of the New York Academy of Sciences 1279, 90–96.

Bardoni, R., Barry, D.M., Li, H., Shen, K.-F., Jeffry, J., Yang, Q., Comitato, A., Li, Y.-Q., and Chen, Z.-F. (2018). Counter-stimuli Inhibit GRPR Neurons via GABAergic Signaling in the Spinal Cord. BioRxiv.

Betley, J.N., Wright, C.V.E., Kawaguchi, Y., Erdélyi, F., Szabó, G., Jessell, T.M., and Kaltschmidt, J.A. (2009). Stringent Specificity in the Construction of a GABAergic Presynaptic Inhibitory Circuit. Cell 139, 161–174.

Borromeo, M.D., Meredith, D.M., Castro, D.S., Chang, J.C., Tung, K.C., Guillemot, F., and Johnson, J.E. (2014). A transcription factor network specifying inhibitory versus excitatory neurons in the dorsal spinal cord. Development 141, 3102–3102.

Bourane, S., Duan, B., Koch, S.C., Dalet, A., Britz, O., Garcia-Campmany, L., Kim, E., Cheng, L., Ghosh, A., Ma, Q., et al. (2015a). Gate control of mechanical itch by a subpopulation of spinal cord interneurons. Science 350, 550–554.

Bourane, S., Grossmann, K.S., Britz, O., Dalet, A., Del Barrio, M.G., Stam, F.J., Garcia-Campmany, L., Koch, S., and Goulding, M. (2015b). Identification of a Spinal Circuit for Light Touch and Fine Motor Control. Cell 160, 503–515.

Bouvier, J., Caggiano, V., Leiras, R., Caldeira, V., Bellardita, C., Balueva, K., Fuchs, A., and Kiehn, O. (2015). Descending Command Neurons in the Brainstem that Halt Locomotion. Cell 163, 1191–1203.

Brenner, D.S., Golden, J.P., and Gereau, R.W. (2012). A Novel Behavioral Assay for Measuring Cold Sensation in Mice. PLoS ONE 7, e39765.

Britz, O., Zhang, J., Grossmann, K.S., Dyck, J., Kim, J.C., Dymecki, S., Gosgnach, S., and Goulding, M. (2015). A genetically defined asymmetry underlies the inhibitory control of flexor– extensor locomotor movements. ELife 4.

Bröhl, D., Strehle, M., Wende, H., Hori, K., Bormuth, I., Nave, K.-A., Müller, T., and Birchmeier, C. (2008). A transcriptional network coordinately determines transmitter and peptidergic fate in the dorsal spinal cord. Developmental Biology 322, 381–393.

Brownstone, R.M., and Chopek, J.W. (2018). Reticulospinal Systems for Tuning Motor Commands. Frontiers in Neural Circuits 12.

Capelli, P., Pivetta, C., Soledad Esposito, M., and Arber, S. (2017). Locomotor speed control circuits in the caudal brainstem. Nature.

Chapman, E. and Tremblay, F. (2015), Scholarpedia, 10(3):7953

Cheng, L., Arata, A., Mizuguchi, R., Qian, Y., Karunaratne, A., Gray, P.A., Arata, S., Shirasawa, S., Bouchard, M., Luo, P., et al. (2004). Tlx3 and Tlx1 are post-mitotic selector genes determining glutamatergic over GABAergic cell fates. Nature Neuroscience 7, 510–517.

Cheng, L., Samad, O.A., Xu, Y., Mizuguchi, R., Luo, P., Shirasawa, S., Goulding, M., and Ma, Q. (2005). Lbx1 and Tlx3 are opposing switches in determining GABAergic versus glutamatergic transmitter phenotypes. Nature Neuroscience 8, 1510–1515.

Comer, J.D., Pan, F.C., Willet, S.G., Haldipur, P., Millen, K.J., Wright, C.V.E., and Kaltschmidt, J.A. (2015). Sensory and spinal inhibitory dorsal midline crossing is independent of Robo3. Frontiers in Neural Circuits 9.

Datta, S.R., Anderson, D.J., Branson, K., Perona, P., and Leifer, A. (2019). Computational Neuroethology: A Call to Action. Neuron 104, 11–24.

Delile, J., Rayon, T., Melchionda, M., Edwards, A., Briscoe, J., and Sagner, A. (2019). Single cell transcriptomics reveals spatial and temporal dynamics of gene expression in the developing mouse spinal cord. Development 146, dev173807.

Detloff, M.R., Fisher, L.C., Deibert, R.J., and Basso, D.M. (2012). Acute and Chronic Tactile Sensory Testing after Spinal Cord Injury in Rats. Journal of Visualized Experiments.

Djouhri, L. (2016). Aδ-fiber low threshold mechanoreceptors innervating mammalian hairy skin: A review of their receptive, electrophysiological and cytochemical properties in relation to Aδ-fiber high threshold mechanoreceptors. Neuroscience & Biobehavioral Reviews 61, 225–238.

Dong, X., and Dong, X. (2018). Peripheral and Central Mechanisms of Itch. Neuron 98, 482–494.

Duan, B., Cheng, L., Bourane, S., Britz, O., Padilla, C., Garcia-Campmany, L., Krashes, M., Knowlton, W., Velasquez, T., Ren, X., et al. (2014). Identification of Spinal Circuits Transmitting and Gating Mechanical Pain. Cell 159, 1417–1432.

Escalante, A., Murillo, B., Morenilla-Palao, C., Klar, A., and Herrera, E. (2013). Zic2-Dependent Axon Midline Avoidance Controls the Formation of Major Ipsilateral Tracts in the CNS. Neuron 80, 1392–1406.

Feng, J., Luo, J., Yang, P., Du, J., Kim, B.S., and Hu, H. (2018). Piezo2 channel-Merkel cell signaling modulates the conversion of touch to itch. Science 360, 530–533.

Fink, A.J.P., Croce, K.R., Huang, Z.J., Abbott, L.F., Jessell, T.M., and Azim, E. (2014). Presynaptic inhibition of spinal sensory feedback ensures smooth movement. Nature 509, 43–48.

Foster, E., Wildner, H., Tudeau, L., Haueter, S., Ralvenius, W.T., Jegen, M., Johannssen, H., Hösli, L., Haenraets, K., Ghanem, A., et al. (2015). Targeted Ablation, Silencing, and Activation Establish Glycinergic Dorsal Horn Neurons as Key Components of a Spinal Gate for Pain and Itch. Neuron 85, 1289–1304.

François, A., Low, S.A., Sypek, E.I., Christensen, A.J., Sotoudeh, C., Beier, K.T., Ramakrishnan, C., Ritola, K.D., Sharif-Naeini, R., Deisseroth, K., et al. (2017). A Brainstem-Spinal Cord Inhibitory Circuit for Mechanical Pain Modulation by GABA and Enkephalins. Neuron 1–41.

Frost, W.N., Tian, L.-M., Hoppe, T.A., Mongeluzi, D.L., and Wang, J. (2003). A cellular mechanism for prepulse inhibition. Neuron 40, 991–1001.

Fukuoka, M., Miyachi, Y., and Ikoma, A. (2013). Mechanically evoked itch in humans: Pain 154, 897–904.

Gao, Z.-R., Chen, W.-Z., Liu, M.-Z., Chen, X.-J., Wan, L., Zhang, X.-Y., Yuan, L., Lin, J.-K.,Wang, M., Zhou, L., et al. (2019). Tac1-Expressing Neurons in the Periaqueductal Gray Facilitate the Itch-Scratching Cycle via Descending Regulation. Neuron 101, 45–59.e9.

Geuther, B.Q., Deats, S.P., Fox, K.J., Murray, S.A., Braun, R.E., White, J.K., Chesler, E.J., Lutz, C.M., and Kumar, V. (2019). Robust mouse tracking in complex environments using neural networks. Communications Biology 2.

Glasgow, S.M. (2005). Ptf1a determines GABAergic over glutamatergic neuronal cell fate in the spinal cord dorsal horn. Development 132, 5461–5469.

Gross, M.K., Dottori, M., and Goulding, M. (2002). Lbx1 specifies somatosensory association interneurons in the dorsal spinal cord. Neuron 34, 535–549.

Hafenreffer, S. (1660). Nosodochion : in quo cutis, eique adhaerentium partium, affectus omnes, singulari methodo et cognoscendi et curandi fidelissime traduntur… (Ulm: Kühn).

Helms, A.W., and Johnson, J.E. (2003). Specification of dorsal spinal cord interneurons. Current Opinion in Neurobiology 13, 42–49.

Hori, K., Cholewa-Waclaw, J., Nakada, Y., Glasgow, S.M., Masui, T., Henke, R.M., Wildner, H., Martarelli, B., Beres, T.M., Epstein, J.A., et al. (2008). A nonclassical bHLH Rbpj transcription factor complex is required for specification of GABAergic neurons independent of Notch signaling. Genes & Development 22, 166–178.

Huang, M., Huang, T., Xiang, Y., Xie, Z., Chen, Y., Yan, R., Xu, J., and Cheng, L. (2008). Ptf1a, Lbx1 and Pax2 coordinate glycinergic and peptidergic transmitter phenotypes in dorsal spinal inhibitory neurons. Developmental Biology 322, 394–405.

Huang, T., Lin, S.-H., Malewicz, N.M., Zhang, Y., Zhang, Y., Goulding, M., LaMotte, R.H., and Ma, Q. (2019). Identifying the pathways required for coping behaviours associated with sustained pain. Nature 565, 86–90.

Hughes, D.I., Mackie, M., Nagy, G.G., Riddell, J.S., Maxwell, D.J., Szabo, G., Erdelyi, F., Veress, G., Szucs, P., Antal, M., et al. (2005). P boutons in lamina IX of the rodent spinal cord express high levels of glutamic acid decarboxylase-65 and originate from cells in deep medial dorsal horn. Proceedings of the National Academy of Sciences 102, 9038–9043.

Ikoma, A., Handwerker, H., Miyachi, Y., and Schmelz, M. (2005). Electrically evoked itch in humans: Pain 113, 148–154.

Jakobsson, J.E.T., Ma, H., and Lagerström, M.C. (2019). Neuropeptide Y in itch regulation. Neuropeptides 78, 101976.

Juravle, G., Binsted, G., and Spence, C. (2017). Tactile suppression in goal-directed movement. Psychonomic Bulletin & Review 24, 1060–1076.

Kabra, M., Robie, A.A., Rivera-Alba, M., Branson, S., and Branson, K. (2013). JAABA: interactive machine learning for automatic annotation of animal behavior. Nature Methods 10, 64–67.

Kawaguchi, Y., Cooper, B., Gannon, M., Ray, M., MacDonald, R.J., and Wright, C.V.E. (2002). The role of the transcriptional regulator Ptf1a in converting intestinal to pancreatic progenitors. Nature Genetics 32, 128–134.

Kini, S.P. (2011). The Impact of Pruritus on Quality of Life: The Skin Equivalent of Pain. Archives of Dermatology 147, 1153.

Lallemend, F., and Ernfors, P. (2012). Molecular interactions underlying the specification of sensory neurons. Trends in Neurosciences 35, 373–381.

Li, Y., Qiu, Q., Watson, S.S., Schweitzer, R., and Johnson, R.L. (2010). Uncoupling skeletal and connective tissue patterning: conditional deletion in cartilage progenitors reveals cell-autonomous requirements for Lmx1b in dorsal-ventral limb patterning. Development 137, 1181–1188.

Liang, H., Watson, C., and Paxinos, G. (2016). Terminations of reticulospinal fibers originating from the gigantocellular reticular formation in the mouse spinal cord. Brain Structure and Function 221, 1623–1633.

Madisen, L., Zwingman, T.A., Sunkin, S.M., Oh, S.W., Zariwala, H.A., Gu, H., Ng, L.L., Palmiter, R.D., Hawrylycz, M.J., Jones, A.R., et al. (2010). A robust and high-throughput Cre reporting and characterization system for the whole mouse brain. Nature Neuroscience 13, 133–140.

Madisen, L., Garner, A.R., Shimaoka, D., Chuong, A.S., Klapoetke, N.C., Li, L., van der Bourg, A., Niino, Y., Egolf, L., Monetti, C., et al. (2015). Transgenic Mice for Intersectional Targeting of Neural Sensors and Effectors with High Specificity and Performance. Neuron 85, 942–958.

Mende, M., Fletcher, E.V., Belluardo, J.L., Pierce, J.P., Bommareddy, P.K., Weinrich, J.A., Kabir, Z.D., Schierberl, K.C., Pagiazitis, J.G., Mendelsohn, A.I., et al. (2016). Sensory-Derived Glutamate Regulates Presynaptic Inhibitory Terminals in Mouse Spinal Cord. Neuron 90, 1189–1202.

Meredith, D.M., Masui, T., Swift, G.H., MacDonald, R.J., and Johnson, J.E. (2009). Multiple Transcriptional Mechanisms Control Ptf1a Levels during Neural Development Including Autoregulation by the PTF1-J Complex. The Journal of Neuroscience: The Official Journal of the Society for Neuroscience 29, 11139–11148.

Mizuguchi, R., Kriks, S., Cordes, R., Gossler, A., Ma, Q., and Goulding, M. (2006). Ascl1 and Gsh1/2 control inhibitory and excitatory cell fate in spinal sensory interneurons. Nature Neuroscience 9, 770–778.

Mollanazar, N.K., Sethi, M., Rodriguez, R.V., Nattkemper, L.A., Ramsey, F.V., Zhao, H., and Yosipovitch, G. (2016). Retrospective analysis of data from an itch center: Integrating validated tools in the electronic health record. Journal of the American Academy of Dermatology 75, 842–844.

Moreno-López, Y., Olivares-Moreno, R., Cordero-Erausquin, M., and Rojas-Piloni, G. (2016). Sensorimotor Integration by Corticospinal System. Frontiers in Neuroanatomy 10.

Mueller, S., Fischer, M., Herger, S., Nüesch, C., Egloff, C., Itin, P., Cajacob, L., Brandt, O., and Mündermann, A. (2019). Good vibrations: Itch induction by whole body vibration exercise without the need of a pruritogen. Experimental Dermatology 28, 1390–1396.

Müller, T., Brohmann, H., Pierani, A., Heppenstall, P.A., Lewin, G.R., Jessell, T.M., and Birchmeier, C. (2002). The homeodomain factor lbx1 distinguishes two major programs of neuronal differentiation in the dorsal spinal cord. Neuron 34, 551–562.

Nakhai, H., Sel, S., Favor, J., Mendoza-Torres, L., Paulsen, F., Duncker, G.I.W., and Schmid, R.M. (2007). Ptf1a is essential for the differentiation of GABAergic and glycinergic amacrine cells and horizontal cells in the mouse retina. Development 134, 1151–1160.

Oetjen, L.K., Mack, M.R., Feng, J., Whelan, T.M., Niu, H., Guo, C.J., Chen, S., Trier, A.M., Xu, A.Z., Tripathi, S.V., et al. (2017). Sensory Neurons Co-opt Classical Immune Signaling Pathways to Mediate Chronic Itch. Cell 171, 217–228.e13.

Orefice, L.L., Zimmerman, A.L., Chirila, A.M., Sleboda, S.J., Head, J.P., and Ginty, D.D. (2016). Peripheral Mechanosensory Neuron Dysfunction Underlies Tactile and Behavioral Deficits in Mouse Models of ASDs. Cell 166, 299–313.

Orefice, L.L., Mosko, J.R., Morency, D.T., Wells, M.F., Tasnim, A., Mozeika, S.M., Ye, M., Chirila, A.M., Emanuel, A.J., Rankin, G., et al. (2019). Targeting Peripheral Somatosensory Neurons to Improve Tactile-Related Phenotypes in ASD Models. Cell 178, 867–886.e24.

Pagani, M., Albisetti, G.W., Sivakumar, N., Wildner, H., Santello, M., Johannssen, H.C., and Zeilhofer, H.U. (2019). How Gastrin-Releasing Peptide Opens the Spinal Gate for Itch. Neuron 103, 102–117.e5.

Paixão, S., Loschek, L., Gaitanos, L., Alcalà Morales, P., Goulding, M., and Klein, R. (2019). Identification of Spinal Neurons Contributing to the Dorsal Column Projection Mediating Fine Touch and Corrective Motor Movements. Neuron.

Pan, H., Fatima, M., Li, A., Lee, H., Cai, W., Horwitz, L., Hor, C.C., Zaher, N., Cin, M., Slade, H., et al. (2019). Identification of a Spinal Circuit for Mechanical and Persistent Spontaneous Itch. Neuron.

Pereira, P.J.S., Machado, G.D.B., Danesi, G.M., Canevese, F.F., Reddy, V.B., Pereira, T.C.B., Bogo, M.R., Cheng, Y.-C., Laedermann, C., Talbot, S., et al. (2015). GRPR/PI3K : Partners in Central Transmission of Itch. Journal of Neuroscience 35, 16272–16281.

Poulet, J., and Hedwig, B. (2006). The cellular basis of a corollary discharge. Science 311, 518–522.

Punnakkal, P., von Schoultz, C., Haenraets, K., Wildner, H., and Zeilhofer, H.U. (2014). Morphological, biophysical and synaptic properties of glutamatergic neurons of the mouse spinal dorsal horn: Glutamatergic neurons of the mouse dorsal horn. The Journal of Physiology 592, 759–776.

Rudomin, P. (2009). In search of lost presynaptic inhibition. Experimental Brain Research 196, 139–151.

Rudomin, P., and Schmidt, R.F. (1999). Presynaptic inhibition in the vertebrate spinal cord revisited. Experimental Brain Research. Experimentelle Hirnforschung. Expérimentation Cérébrale 129, 1–37.

Schut, C., Dalgard, F., Halvorsen, J., Gieler, U., Lien, L., Aragones, L., Poot, F., Jemec, G., Misery, L., Kemény, L., et al. (2019). Occurrence, Chronicity and Intensity of Itch in a Clinical Consecutive Sample of Patients with Skin Diseases: A Multi-centre Study in 13 European Countries. Acta Dermato Venereologica 99, 146–151.

Sciolino, N.R., Plummer, N.W., Chen, Y.-W., Alexander, G.M., Robertson, S.D., Dudek, S.M., McElligott, Z.A., and Jensen, P. (2016). Recombinase-Dependent Mouse Lines for Chemogenetic Activation of Genetically Defined Cell Types. Cell Reports 15, 2563–2573.

Seki, K., Perlmutter, S.I., and Fetz, E.E. (2003). Sensory input to primate spinal cord is presynaptically inhibited during voluntary movement. Nature Neuroscience 6, 1309–1316.

Sorkin, L.S., and Puig, S. (1996). Neuronal model of tactile allodynia produced by spinal strychnine: effects of excitatory amino acid receptor antagonists and a mu-opiate receptor agonist. Pain 68, 283–292.

Sorkin, L.S., Puig, S., and Jones, D.L. (1998). Spinal bicuculline produces hypersensitivity of dorsal horn neurons: effects of excitatory amino acid antagonists. Pain 77, 181–190.

Sukhtankar, D.D., and Ko, M.-C. (2013). Physiological Function of Gastrin-Releasing Peptide and Neuromedin B Receptors in Regulating Itch Scratching Behavior in the Spinal Cord of Mice. PLoS ONE 8, e67422.

Sun, Y.-G., and Chen, Z.-F. (2007). A gastrin-releasing peptide receptor mediates the itch sensation in the spinal cord. Nature 448, 700–703.

Sun, Y.-G., Zhao, Z.-Q., Meng, X.-L., Yin, J., Liu, X.-Y., and Chen, Z.-F. (2009). Cellular Basis of Itch Sensation. Science 325, 1531–1534.

Takatoh, J., Nelson, A., Zhou, X., Bolton, M.M., Ehlers, M.D., Arenkiel, B.R., Mooney, R., and Wang, F. (2013). New Modules Are Added to Vibrissal Premotor Circuitry with the Emergence of Exploratory Whisking. Neuron 77, 346–360.

Tripathi, R., Knusel, K.D., Ezaldein, H.H., Bordeaux, J.S., and Scott, J.F. (2019). The cost of an itch: A nationally representative retrospective cohort study of pruritus-associated health care expenditure in the United States. Journal of the American Academy of Dermatology 80, 810–813.

Tripodi, M., Stepien, A.E., and Arber, S. (2011). Motor antagonism exposed by spatial segregation and timing of neurogenesis. Nature 479, 61–66.

Ueno, M., Nakamura, Y., Li, J., Gu, Z., Niehaus, J., Maezawa, M., Crone, S.A., Goulding, M., Baccei, M.L., and Yoshida, Y. (2018). Corticospinal Circuits from the Sensory and Motor Cortices Differentially Regulate Skilled Movements through Distinct Spinal Interneurons. Cell Reports 23, 1286–1300.e7.

Usoskin, D., Furlan, A., Islam, S., Abdo, H., Lönnerberg, P., Lou, D., Hjerling-Leffler, J., Haeggström, J., Kharchenko, O., Kharchenko, P.V., et al. (2015). Unbiased classification of sensory neuron types by large-scale single-cell RNA sequencing. Nature Neuroscience 18, 145–153.

Voudouris, D., and Fiehler, K. (2017). Enhancement and suppression of tactile signals during reaching. Journal of Experimental Psychology: Human Perception and Performance 43, 1238–1248.

Vrontou, S., Wong, A.M., Rau, K.K., Koerber, H.R., and Anderson, D.J. (2013). Genetic identification of C fibres that detect massage-like stroking of hairy skin in vivo. Nature 493, 669–673.

Wahlgren, C.F., Hagermark, O., and Bergstrom, R. (1991). Patients’ perception of itch induced by histamine, compound 48/80 and wool fibres in atopic dermatitis. Acta Dermato Venereologica 71, 488–494.

Watson, A.H. (1992). Presynaptic modulation of sensory afferents in the invertebrate and vertebrate nervous system. Comparative Biochemistry and Physiology. Comparative Physiology 103, 227–239.

Weisshaar, E., Szepietowski, J., Dalgard, F., Garcovich, S., Gieler, U., Giménez-Arnau, A., Lambert, J., Leslie, T., Mettang, T., Misery, L., et al. (2019). European S2k Guideline on Chronic Pruritus. Acta Dermato Venereologica 99, 469–506.

Wickersham, I.R., Finke, S., Conzelmann, K.-K., and Callaway, E.M. (2006). Retrograde neuronal tracing with a deletion-mutant rabies virus. Nature Methods 4, 47–49.

Wildner, H. (2006). dILA neurons in the dorsal spinal cord are the product of terminal and non-terminal asymmetric progenitor cell divisions, and require Mash1 for their development. Development 133, 2105–2113.

Wildner, H., Das Gupta, R., Brohl, D., Heppenstall, P.A., Zeilhofer, H.U., and Birchmeier, C. (2013). Genome-Wide Expression Analysis of Ptf1a- and Ascl1-Deficient Mice Reveals New Markers for Distinct Dorsal Horn Interneuron Populations Contributing to Nociceptive Reflex Plasticity. Journal of Neuroscience 33, 7299–7307.

Yosipovitch, G., and Bernhard, J.D. (2013). Chronic Pruritus. New England Journal of Medicine 368, 1625–1634.

Zeilhofer, H.U., Wildner, H., and Yévenes, G.E. (2012). Fast Synaptic Inhibition in Spinal Sensory Processing and Pain Control. Physiological Reviews 92, 193–235.

Zimmerman, A.L., Kovatsis, E.M., Pozsgai, R.Y., Tasnim, A., Zhang, Q., and Ginty, D.D. (2019). Distinct Modes of Presynaptic Inhibition of Cutaneous Afferents and Their Functions in Behavior. Neuron 102, 420–434.e8.

